# RNA scaffolds the Golgi ribbon by forming condensates with GM130

**DOI:** 10.1101/2023.03.02.530905

**Authors:** Yijiun Zhang, Joachim Seemann

## Abstract

The mammalian Golgi apparatus is composed of stacks of cisternae that are laterally linked to form a continuous ribbon like structure, but the molecular mechanisms that maintain the Golgi ribbon remain unclear. Here, we show that ribbon formation is mediated by biomolecular condensates of RNA and the Golgi resident protein GM130. We identified GM130 as a membrane-bound RNA binding protein at the Golgi. Acute degradation of either RNA or GM130 in cells disrupted the Golgi ribbon. Under stress conditions, RNA was displaced from GM130 and the ribbon was disjoint, which was restored after cells recovered from stress. When overexpressed in cells, GM130 formed RNA-dependent liquid-like condensates. GM130 contains an intrinsically disordered domain at its N-terminus, which was sufficient to recruit RNA to drive condensate assembly *in vitro*. Condensates of the N-terminal domain of GM130 and RNA were sufficient to link purified rat liver Golgi membranes which is a reconstruction of aspects of lateral linking of stacks into a ribbon-like structure. Together, these studies reveal that biomolecular condensates of GM130-RNA scaffold the Golgi ribbon.

## Introduction

The Golgi apparatus is a membrane-bound organelle required for post-translational processing of secreted proteins. It receives proteins and lipids from the endoplasmic reticulum and sorts them to their destinations such as endosomes, lysosomes or the plasma membrane (Pantazopoulou and Glick, 2019). Its function as the central sorting and modification hub of the secretory pathway is conserved among eukaryotes and facilitated by stacks of disk-shaped membrane cisternae. Lower eukaryotes such as worms or flies, contain a low number of stacks, about 25 in flies, that are scattered throughout the cytoplasm (Kondylis et al., 2007). In mammalian cells the Golgi is comprised of 100-200 stacks, but in contrast to non-vertebrate cells, the stacks are laterally fused together by tubular membranes into a twisted contiguous structure referred to as the Golgi ribbon (Nakamura et al., 2012). This higher-order organization into a joint compartment allows the Golgi to accommodate large proteins that are bigger than individual cisterna such as collagen or Weibel-Palade bodies (Lavieu et al., 2014; Page et al., 2022).

In most cell types, the Golgi ribbon localizes next to the centrosomes in the perinuclear region of the cell. This focused central concentration of the ribbon defines a polarity axis to direct vesicles sorting to a specific region of the plasma membrane, which is important for cell polarization and differentiation (Wei and Seemann, 2017; Mascanzoni et al., 2022). On the other hand, the integrity of the Golgi ribbon is perturbed in cells exposed to stress conditions causing Golgi dysfunction (Machamer, 2015). A compromised ribbon is commonly associated with a broad array of diseases, including neurodegenerative diseases, muscular dystrophies (Zappa et al., 2018; Li et al., 2019). Golgi fragmentation is a very early event preceding disease phenotypes in ALS or Alzheimer’s disease (van Dis et al., 2014; Haukedal et al., 2022), suggesting that Golgi fragmentation may cause disease onset, but the underlying mechanisms are largely unclear.

The unique structure of the Golgi ribbon is highly dynamic and constantly remodeled by budding and fusion of transport vesicles, yet the characteristic ribbon structure is maintained (Lowe, 2011). The ribbon is thought to be stabilized by multimeric interactions of structural Golgi resident proteins that collectively form a matrix (Slusarewicz et al., 1994; Seemann et al., 2000). The Golgi matrix was initially visualized by EM as proteinaceous 11 nm projections surrounding the cisternae (Cluett and Brown, 1992). Golgi matrix proteins are foremost proteins of the Golgin and GRASP families. Golgins are long rod-like proteins with extensive coiled-coil domains that are attached to the cytoplasmic face of the Golgi having the potential to support long-range interactions (Gillingham and Munro, 2016). The matrix components define the higher order organization of the Golgi ribbon and are necessary and sufficient to build and maintain the ribbon structure in the perinuclear region (Seemann et al., 2000). Furthermore, when the Golgi is removed from cells by laser ablation, a new Golgi can reform within 12 hours with Golgi matrix proteins appearing first followed by other Golgi components. Knockdown of the Golgin GM130 suppressed Golgi biogenesis (Tängemo et al., 2011), indicating that GM130 is a central component of that organizes the matrix.

Knockdown of GRASP55 or GRASP65 by RNAi caused Golgi fragmentation and ribbon unlinking, suggesting that GRASP proteins are organizing the lateral connections between adjacent cisternae to link stacks into a ribbon (Sütterlin et al., 2005; Duran et al., 2008; Feinstein and Linstedt, 2008; Puthenveedu et al., 2006). In addition, GM130 was also implicated in the process as it recruits GRASP65 to the Golgi to from a complex (Puthenveedu et al., 2006). However, acute degradation of GRASP55 or GRASP65 within 2 h did not affect the lateral connections of the ribbon, while depletion of both GRASPs indirectly caused a partial loss of GM130 and other Golgins thereby affecting ribbon integrity (Zhang and Seemann, 2021). This suggests that GRASPs are indirectly involved in linking the ribbon and other unidentified molecules may directly function as lining factors.

In this study, we set out to define the molecules that are required to link the Golgi ribbon. We identified GM130 as a Golgi-localized RNA binding protein, which recruits RNA and associated RNA binding proteins to Golgi membranes. Acute degradation of either RNA or GM130 in cells disrupts the ribbon without affecting the organization of stacks. Under stress conditions, RNA disassociates from GM130 while the Golgi ribbon is disrupted and fragmented. Furthermore, RNA binding of GM130 induces liquid-liquid phase separation, which is sufficient to link purified Golgi membranes together. Our results show that RNA functions as a structural element that scaffolds the Golgi ribbon by a forming a biomolecular condensate with GM130.

## Results

### GM130 forms a complex with RNA-binding proteins

To identify molecules that specifically associate with the GM130/GRASP65 complex relative to Golgin-45/GRASP55, we used our previously established RC55 and RC65 cell lines (Zhang and Seemann, 2021). Both cell lines are gene edited SV589 human fibroblasts expressing GRASP55 (RC55) or GRASP65 (RC65) endogenously tagged with a Flag epitope and an auxin-inducible degron (AID) sequence (Figure 1A). Lysates from RC65 and RC55 cells were incubated with anti-Flag beads, bound complexes were eluted with a Flag peptide and identified by semi-quantitative mass spectrometry. We then plotted the relative abundance of the detected proteins to identify components that were preferentially enriched with GRASP65 compared with GRASP55 (Figure 1B). To validate this approach, we used Golgin-45 and GM130, which form distinct complexes with GRASP55 and GRASP65, respectively (Barr et al., 1998; Short et al., 2001). As expected, GRASP55 preferentially pulled down Golgin-45, whereas GRASP65 significantly enriched GM130. As in previous reports, we also found several members of the p24/transmembrane emp24 domain family of cargo receptors (TMED) in a complex with GRASP55 (Barr et al., 2001; Ahat et al., 2022). Strikingly, mass spectrometry analysis further identified several RNA-binding proteins (RBPs) that specifically associate with GRASP65, including G3BP1 (G3BP Stress Granule Assembly Factor 1), FXR1 (Fragile X mental retardation syndrome-related protein 1), CAPRIN1 (also known as RNA granule protein 105 or RNG105), and PABPC1 (PABP/Polyadenylate-binding protein 1). Western blotting analysis confirmed that GM130, FXR1, G3BP1 and PABC1 co-immunoprecipitated with GRASP65-Flag, but not with GRASP55-Flag (Figure 1C). Furthermore, FXR1 and GM130 were also co-pelleted with endogenous GRASP65 from SV589 cell lysates using anti-GRASP65 antibodies (Figure 1D).

**Figure 1.**
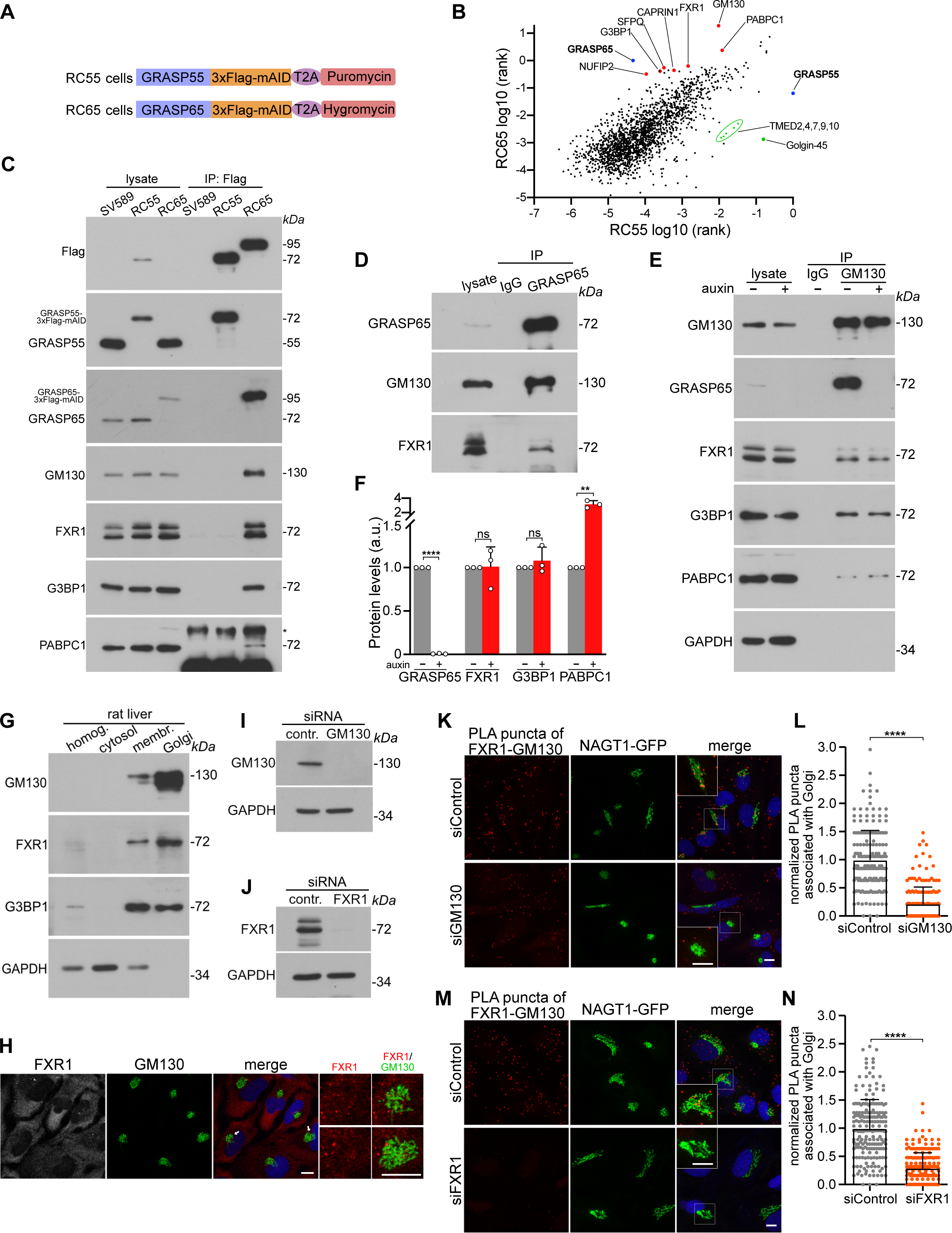
**GM130 is in a complex with the RNA binding proteins FXR1, G3BP1 and PABPC1** (A). Scheme of endogenous GRASP55 or GRASP65 tagged with 3xFlag and mAID (minimal auxin inducible degron) sequence followed by a self-cleaving T2A peptide, puromycin or hygromycin selection gene. RC55 cells express endogenous GRASP55-3xFlag-mAID, RC65 cells express GRASP65-3xFlag-mAID. (B). Scatterplot showing of proteins enriched with GRASP65 compared to GRASP55 identified by mass spectrometry. Flag-tagged GRASP55 and GRASP65 were immunoprecipitated from RC55 or RC65 cell lysates and associated proteins were analyzed by mass spectrometry. The abundance of each interactor was normalized to GRASP55 or GRASP65 and plotted by rank. GRASP55 and GRASP65 (blue), representative interactors of GRASP55 (green), representative interactors of GRASP65 (orange) are highlighted. The results are from one of two independent experiments showing comparable results. (C). Verification of the mass spectrometry results by immunoprecipitating Flag from the lysates of wild-type SV589, RC55 and RC65 cells and immunoblotting with the indicated antibodies. IP, immunoprecipitation. * denotes unspecific band. (D). FXR1 associates with the GRASP65-GM130 complex. Endogenous GRASP65 was immunoprecipitated from SV589 cell lysates and analyzed by western blotting. (E). GM130 associates with FXR1, G3BP1 and PABPC1 independently of GRASP65. RC65 cells were treated with doxycycline for 6 h to express the E3 ubiquitin ligase component TIR1and then with the auxin IAA for another 2 h to degrade GRASP65. GM130 was immunoprecipitated from the cell lysates and subjected to Western blotting with the indicated antibodies. (F). Quantitation of (E). The intensities of each protein were normalized to controls. n = 3 independent experiments. ** P < 0.01; **** P < 0.0001; ns, not significant. Error bars represent mean ± SD. (G). FXR1 and G3BP1 are enriched on purified rat liver Golgi membranes. Immunoblots of 10 µg protein of rat liver homogenate (homog.), cytosol, membrane-enriched intermediate purification fraction (membr.) and purified Golgi membranes. (H). FXR1 foci associate with GM130. SV589 cells were immunostained for GM130 together with FXR1. DNA is labelled in blue. The Golgi areas marked by arrows are shown magnified in the right panels. Scale bars, 10 µm. (I-M). FXR1 association with GM130 visualized by proximity ligation assay (PLA) in SV589 cells. PLA of FXR1-GM130 (K, M) in SV589 cells expressing the Golgi marker NAGTI-GFP transfected with control siRNA or siRNA against GM130. PLA puncta, red; DNA, blue. The Golgi area marked by a box was magnified in the inset. Scale bars, 10 µm. Quantitation of the number of PLA puncta associated with the Golgi marked by NAGTI-GFP is shown in (L) n = 2 independent experiments, **** P < 0.0001 and in (N) n = 3 independent experiments, **** P < 0.0001. Immunoblots of cells transfected with siRNA are shown in (I) and (J).

Given that GRASP65 forms a tight complex with GM130 via its PDZ domain on Golgi membranes (Barr et al., 1998), we tested whether the identified RNA-binding proteins associate with GRASP65 or GM130. For this purpose, we used RC65 cells that express the E3 ubiquitin ligase component TIR1 and endogenous GRASP65 tagged with an AID sequence. TIR1 expression was first induced for 6 h before cells were treated with auxin for 2 h to degrade GRASP65 by the proteasome. Consistent with our previous report (Zhang and Seemann, 2021), GRASP65 was efficiently depleted without affecting GM130 levels (Figure 1E). Notably, FXR1, G3BP1 and PABC1 associated with immunoprecipitated endogenous GM130 whether GRASP65 was present or degraded (Figure 1 E, F). Thus, the RBPs associate with GM130 independently of GRASP65.

### The complex of GM130 and RBPs is localized on Golgi membranes

The localization of GM130 is restricted to the Golgi apparatus (Nakamura et al., 1995), while FXR1 and G3BP1 were characterized as primarily cytosolic proteins (Siomi et al., 1995; Gallouzi et al., 1998). By associating with GM130, FXR1 should be at least partially present on Golgi membranes. To this end, we used subcellular fractionation to purify Golgi membranes from rat liver (Wang et al., 2006; Wei and Seemann, 2009b). Western blot analysis found FXR1 and G3BP1 to be enriched on Golgi membranes compared to homogenate, cytosol and the membrane-enriched intermediate purification fraction (Figure 1G). Next, we used immunofluorescence microscopy to confirm the presence of FXR1 on the Golgi. FXR1 was distributed throughout the cytoplasm and also concentrated in multiple small granular structures reminiscent of RNA stress granules (Tourrière et al., 2003). These granules tended to accumulate close to the GM130 signal in the prenuclear Golgi region (Figure 1H).

To further assess the association of FXR1 with GM130 on the Golgi, we used an *in-situ* proximity ligation assay (PLA) that can detect protein-protein associations with single molecule resolution as discrete fluorescent foci (Söderberg et al., 2006). We performed PLA with antibodies against FXR1 and GM130 in cells expressing the Golgi marker NAGTI-GFP. As control for the signal specificity, we depleted GM130 or FXR1 by RNAi (Figure 1I, J). In cells treated with control siRNA, PLA puncta were concentrated at the GFP-marked Golgi, thus visualizing association of FXR1 with GM130 (Figure 1 K, M). The number of PLA dots in the Golgi area were significantly diminished in cells depleted of either FXR1 or GM130 (Figure 1 L, N). Taken together, these data indicate that FXR1 and GM130 are present in a complex that localizes to Golgi membranes.

### The association of GM130 with RBPs is RNA dependent

Considering that FXR1, G3BP1 and PABPC1 bind RNA (Riggs et al., 2020), we tested whether their association with GM130 requires RNA. Treatment of GM130 immunoprecipitates with RNase A displaced FXR1, G3BP1 and PABPC1, indicating that RNA is an integral component of their complex with GM130 (Figure 2 A, B). The presence of RNA was further confirmed by extraction of the RNA from the IP complexes followed by agarose gel electrophoresis (Figure 2 C).

**Fig. 2.**
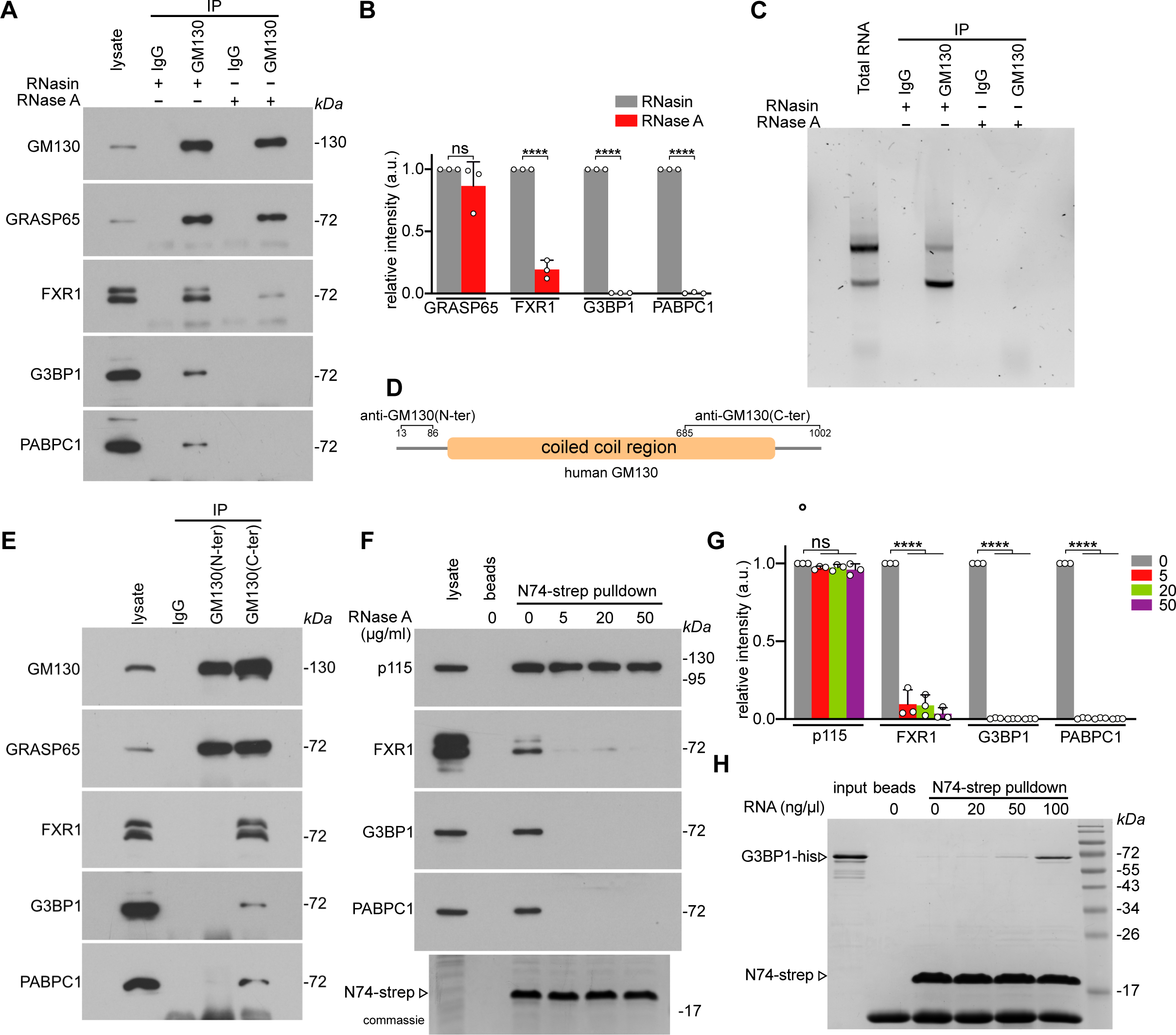
**The association of the RNA-binding proteins FXR1, G3BP1 and PABPC1 with the N-terminal domain of GM130 is RNA dependent** (A). RNase treatment dissociates GM130 from FXR1, G3BP1 and PABPC1. GM130 was immunoprecipitated from SV589 cell lysates and treated with RNase inhibitor (RNasin, 40 U/ml) or RNase A (20 µg/ml) and then subjected to Western blotting. (B). Quantitation of (A). The intensity of each protein was normalized to RNasin-treated sample. n = 3 independent experiments. **** P < 0.0001; ns, not significant. Error bars represent mean ± SD. (C) GM130 binds RNA. GM130 was immunoprecipitated and treated as in (A). RNA was extracted from the samples and separated on an agarose gel. 250 ng total RNA from SV589 cells loaded as control, 250 ng RNA from RNasin-treated samples and equal volume of RNA from other conditions were loaded. (D). Scheme of the human GM130 depicting the epitopes of antibodies against the N-terminal domain of GM130 (anti-GM130(N-ter)) and the C-terminal epitope of anti-GM130(C-ter). (E). Antibodies against the N-terminal domain of GM130 displace RBPs. GM130 was immunoprecipitated from SV589 cell lysates with anti-GM130(N-ter) or anti-GM130(C-ter) and analyzed by immunoblotting. (F). The N-terminal peptide or GM130 (N74) is sufficient to form an RNA-dependent complex with FXR1, G3BP1 and PABPC1. N74-strep was incubated with SV589 cell lysates, pulled down, treated with different concentrations of RNase A and then analyzed by immunoblotting and Coomassie blue staining. (G). Quantitation of (F). The intensity of each protein was normalized to control-treatment. n = 3 independent experiments. **** P < 0.0001; ns, not significant. Error bars represent mean ± SD. (H). RNA mediates the binding of N74 and G3BP1. N74-strep was incubated with G3BP1-his and different concentrations of total RNA. Then N74-strep was pulled down and analyzed by SDS-PAGE and Coomassie blue staining. G3BP1 was only effectively pulled down with increasing RNA concentrations.

To gain insight into the domain of GM130 that associates with RNA and RBPs, we immunoprecipitated GM130 with antibodies raised against the N-terminal domain (N74) (Wei et al., 2015) or with antibodies targeting the C-terminal region (Figure 2 D). Both antibodies pulled down GM130 in complex with GRASP65 as expected. However, C-terminal antibodies also co-pelleted FXR1, G3BP1 and PABPC1, while the N-terminal antibodies did not (Figure 2 E). This result indicates that the antibodies against the N-terminal domain displaced the RBPs. We therefore tested if the N-terminal peptide (N74) is sufficient to form an RNA-dependent complex with RBPs. Lysates of SV589 cells were incubated with N74 linked to a strep tag (N74-strep) followed by pulldown with Strep-tag beads. As a positive control, N74-strep recruited p115, previously reported to bind the N-terminal domain of GM130 (Nakamura et al., 1997). N74 also pulled down FXR1, G3BP1 and PABPC1 and their binding was diminished by RNase treatment of the beads while p115 binding was unaffected (Figure 2 F, G). To directly test whether the complex is RNA mediated, we incubated recombinant G3BP1-his with N74-strep and increasing amounts of total RNA isolated from SV589 cells. N74 pulled down increasing amounts of G3BP1 with rising RNA concentrations (Figure 2 H). Taken together, the data in Figure 2 show that the complex of RBPs with the N-terminal domain of GM130 is RNA-dependent.

### Acute degradation of either RNA or GM130 in cells unlinks the Golgi ribbon

Previous studies relying on chronic downregulation by RNAi implicated GM30 by binding to GRASP65 in lateral joining adjacent Golgi stacks to equilibrate the distribution of Golgi-resident enzymes within the interconnected Golgi ribbon (Puthenveedu et al., 2006). We therefore examined if acute degradation of GM130 or RNA in cells directly affects Golgi ribbon integrity. Microinjection of RNase A into the cytoplasm of SV589 cells disrupted the Golgi ribbon within 1 h, but not in control-injected cells (Figure 3 A). The area occupied by the Golgi increased significantly and the ribbon became disjoint (Figure 3 B, C), but the stacks remained polarized with cis cisternae labelled by GM130 attached to the trans-Golgi marker Golgin-97 (Figure 3 D, E, F). Thus, acute RNA depletion in cells disrupts the lateral association of stacks within the ribbon but does not affect stacking of the cisternae.

**Fig.3.**
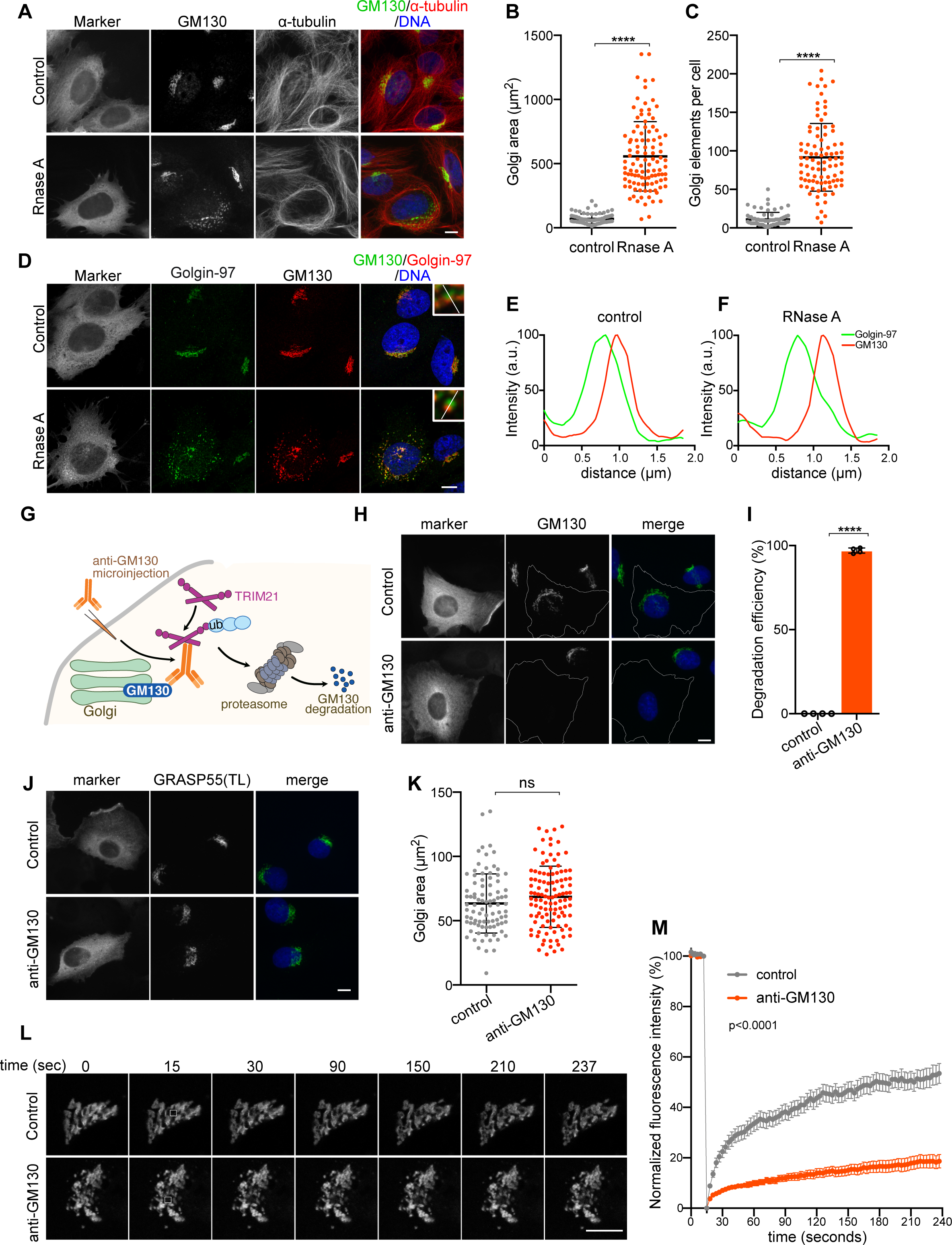
Acute degradation of either RNA or GM130 disrupts the lateral linking of the Golgi ribbon. (A). Microinjection of RNase A fragments the Golgi in 1 h. SV589 cells were injected with 1 mg/ml RNase A together with fluorescent dextran as an injection marker or fluorescent dextran only as control. The cells were fixed 1 hour later and immunostained for GM130 (green) and α-tubulin (red) and labeled for DNA (blue). Scale bar, 10 µm. (B). Quantitation of the Golgi area in control-injected (n = 79) and RNase A-injected (n = 105) SV589 cells from three independent experiments. **** P < 0.0001; Error bars represent mean ± SD. (C). Quantitation of the Golgi elements in control-injected (n = 72) and RNase A-injected (n = 91) SV589 cells from three independent experiments. **** P < 0.0001; Error bars represent mean ± SD. (D-F). Microinjection of RNase A does not affect Golgi stacking. SV589 cells were injected with RNase A or control dextran as in (A). The cells were fixed after 1 h and stained for the trans Golgi protein Golgin-97 (green) and cis-Golgi localized GM130 (red) and DNA (blue). Insets show a magnified Golgi stack with the white line indicating the line scan of the fluorescence intensities of control in (E), and RNase A in (F). Scale bar, 10 µm. (G) Schematic illustration of the Trim-Away method to acutely degrade endogenous GM130. (H-I) Microinjection of anti-GM130 (morsel) acutely degraded GM130 in 2 h. NRK cells stably expressing the ubiquitin ligase mCherry-TRIM21 were injected with anti-GM130 IgG or control IgG (anti-Flag) together with fluorescent dextran as an injection marker. Cells were fixed 2 h later and stained for GM130 (green) and DNA (blue). The degradation efficiency of GM130 was quantified in (H) from four independent experiments with >50 cells analyzed per experiment per condition. **** P < 0.0001. Error bars represent mean ± SD. (J) Acute degradation of GM130 does not increase the Golgi area. Cells from (H) were stained for GRASP55 (green) and DNA (blue). (K) Quantitation of the Golgi area marked by GRASP55 from (J). n = 3 independent experiments with 90 cells from control, 111 cells from morsel analyzed. ns, not significant. Error bars represent mean ± SD. (L) Acute degradation of GM130 causes Golgi fragmentation. NRK cells stably expressing mCherry-TRIM21 and the Golgi marker NAGTI-GFP were injected with anti-GM130 IgG or anti-Flag IgG as control together with fluorescent dextran as an injection marker. FRAP analysis of NAGTI-GFP was then performed in injected cells. Representative images at the indicated time points are shown. White boxes indicate the photobleached area of the Golgi. Scale bar, 10 µm. (M) Quantitation of the FRAP results. The recovery rate at each time point was calculated as the ratio of the mean intensity of the photobleached area to that of the adjacent area and then normalized to the closest time point before bleaching. n = 31 (control) and n = 39 (anti-GM130) cells from four independent experiments. Error bars represent mean ± SEM.

Conversely, since we found that GM130 binds RNA, acute elimination of GM130 should phenocopy RNA depletion, and affect the integrity of the Golgi ribbon. To test this possibility, we acutely degraded GM130 by employing the Trim Away approach (Clift et al., 2017). Trim Away has the advantage of being able to eliminate endogenous proteins without prior modifications. For this approach, IgG binding to the target protein is introduced into cells. The antigen/IgG complex is then recognized by the ubiquitin ligase TRIM21, which directs the complex for rapid degradation by the proteasome (Figure 3 G). Microinjection of anti-GM130 IgG into NRK cells stably expressing TRIM21 resulted in efficient depletion of GM130 within 2 h, while control IgG had no such effect (Figure 3 H, I). The Golgi integrity as marked by GRASP55 appeared altered (Figure 3 J), although the Golgi area was not significantly enlarged (Figure 3 K).

To further assess the effect of GM130 elimination on the integrity of the Golgi ribbon, we expressed the Golgi enzyme NAGTI-GFP and then performed FRAP to analyze GFP recovery as a measure of lateral mobility of NAGTI-GFP within the ribbon (Zhang and Seemann, 2021). Acute elimination of GM130 significantly diminished the recovery of the GFP signal, indicating a loss of connectivity between stacks (Figure 3 L, M). Comparable to RNase injection (Figure 3 D-F), the Golgin-84 labeled cis Golgi remained next to the trans Golgi marked by TGN38, showing that GM130 depletion did not affected stacking (Figure S1 A, B, C). Taken together, these results show that acute disruption of the complex of GM130 and RNA by degradation of either GM130 or RNA severs the Golgi ribbon without affecting stacking.

### Oxidative stress displaces RNA from GM130 and causes Golgi fragmentation

The integrity of the Golgi ribbon is commonly perturbed in cells exposed to stress conditions, including pharmacological or oxidative stress (Machamer, 2015). Because the Golgi ribbon was compromised following disruption of the RNA-GM130 complex, we investigated whether disconnection of the ribbon in cells exposed to stress coincides with RNA displacement from GM130. For this purpose, we treated SV589 cells for 1 h with the commonly used oxidative stress-inducing reagent sodium arsenite. Arsenite triggers the formation of cytoplasmic RNA stress granules (Kedersha et al., 2005), which assemble by liquid-liquid phase separation of mRNA along with RNA-binding proteins including G3BP1, FXR1 and PABPC1 (Riggs et al., 2020; Sanders et al., 2020). We first examined the extent to which stress changes the Golgi structure. Arsenite triggered redistribution of FXR1 from the cytosol into distinct cytoplasmic granules (Figure 4 A), which were dissolved after arsenite was removed. Arsenite-induced stress further resulted in a disrupted the Golgi structure (Figure 4 A). The area occupied by the Golgi was significantly enlarged (Figure 4 B) and the Golgi ribbon was fragmented (Figure 4 C), both of which recovered after arsenite was washed out. FRAP analysis of stressed SV589 cells expressing NAGTI-GFP showed significantly reduced interconnection of the ribbon. Removal of arsenite fully restored the recovery of the GFP signal to control conditions showing that an interconnected Golgi ribbon reformed (Figure 4 D, E).

**Fig.4.**
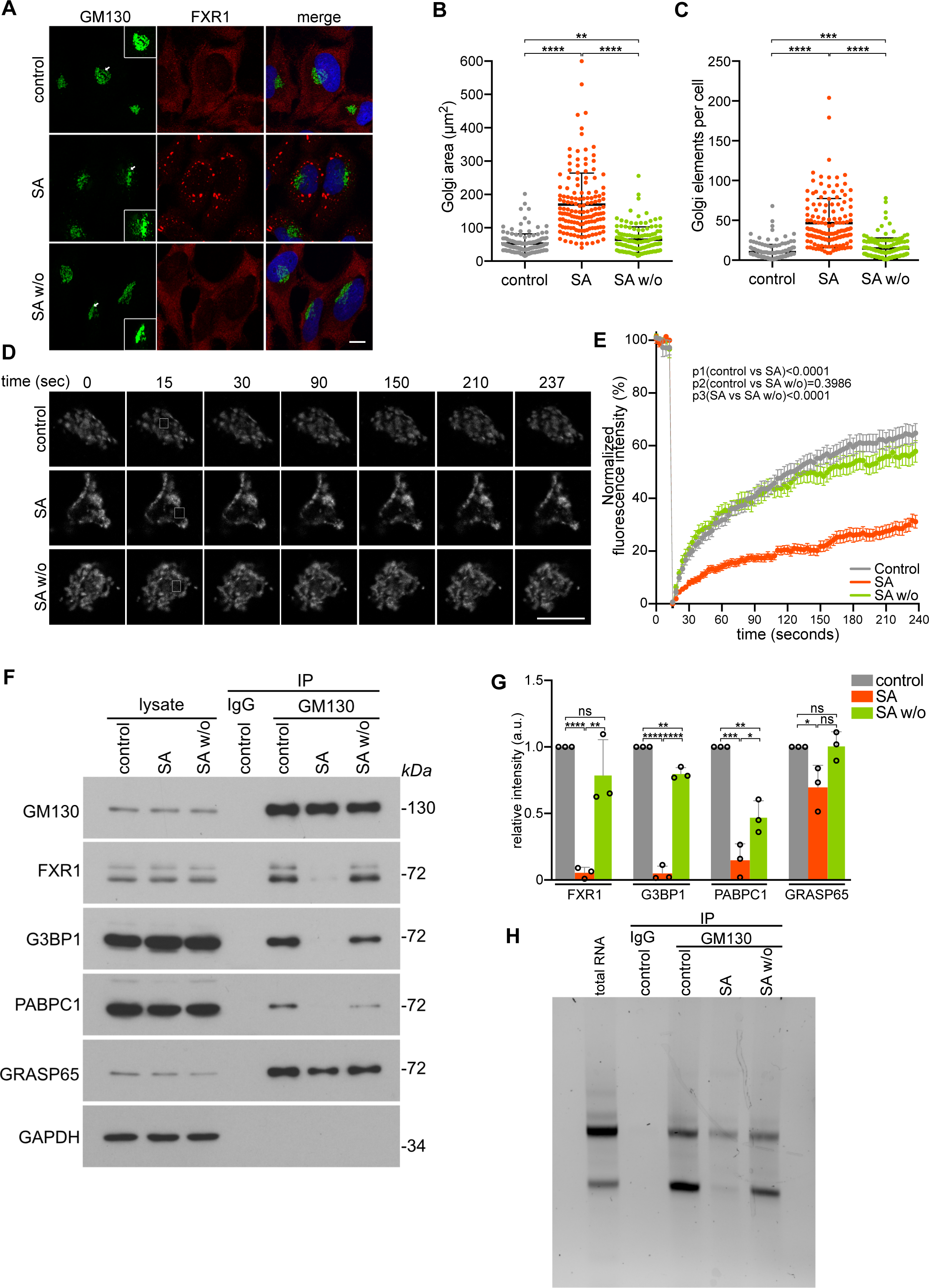
Oxidative stress displaces RNA from GM130 and causes Golgi fragmentation. (A). Sodium arsenite (SA) treatment caused stress granule formation and Golgi fragmentation. SV589 cells were treated with 250 µM SA or were mock treated for 1 h. SA was then washed out and cells were incubated for additional 16 h before fixation (SA w/o). The cells were fixed and immunolabeled for GM130 (green), FXR1 (red) and stained for DNA (blue). Arrows point at the Golgi that is magnified in the insets with brightness increased to visualize Golgi fragments. Scale bar, 10 µm. (B). Quantitation of the Golgi area labeled by GM130 from (A). n = 3 independent experiments with the total number cells analyzed as follows: control: n = 152 cells, SA: n = 148 cells, SA w/o: n = 139 cells. (C). Quantitation of the number of Golgi elements per cell marked by GM130 from (A). n = 3 independent experiments with the total number cells analyzed as follows: control: n = 152 cells, SA: n = 148 cells, SA w/o: n = 139 cells. (D). FRAP analysis of SV589 cells expressing NAGTI-GFP treated with 250 µM SA or mock treated for 1 h. SA was then washed out and cells were then subjected to FRAP analysis (SA) or incubated for 16 h before FRAP analysis (SA w/o). Representative images at the indicated time points are shown, with white boxes indicating the photobleached area of the Golgi. Scale bar, 10 µm. (E). Quantitation of the FRAP results. The recovery rate at each time point was calculated as the ratio of the mean intensity of the photobleached area to that of the adjacent area and then normalized to the closest time point before bleaching. n = 3 independent experiments with the total number cells analyzed as follows: control: n = 30 cells, SA: n = 29cells, SA w/o: n = 30 cells. Error bars represent mean ± SEM. (F). SA treatment causes the loss of FXR1, G3BP1 and PABPC1 from GM130, which is reversed after SA washout. SV589 cells were treated with 250 µM SA (SA) or mock (control) for 1 h. SA was then washed out and cells were incubated for additional 16 h before lysis (SA w/o). GM130 was immunoprecipitated from the cell lysates and separated by immunoblotting with the indicated antibodies. (G). Quantitation of (F). The intensity of each protein was normalized to control conditions. n = 3 independent experiments. * P < 0.05; ** P < 0.01; *** P < 0.001; **** P < 0.0001; ns, not significant. Error bars represent mean ± SD. (H). SA treatment causes the loss of RNA from GM130 but is reversed after SA washout. RNA was extracted from immunoprecipitates from (F) and separated on an agarose gel.

Because loss of RNA or GM130 results in a disjoint Golgi ribbon (Figure 3), we determined whether stress causes dissociation of RNA from GM130. In arsenite-treated cells, association of FXR1, G3BP1 and PABC1 with the GM130-GRASP65 complex was abolished. Consistent with the reforming Golgi integrity, arsenite removal let to reassociation of GM130 with RBPs (Figure 4 F, G). Furthermore, RNA was not extracted from GM130 immunoprecipitates from stressed cells, but we detected RNA after arsenite was washed out (Figure 4 H). Together, these results suggest that the Golgi integrity is compromised under stress conditions due to disruption of RNA binding to GM130.

### GM130 forms RNA-dependent liquid-like condensates in cells

RNA-binding proteins in association with RNA have the propensity to undergo liquid-liquid phase separation to form biomolecular condensates (Lin et al., 2015; Roden and Gladfelter, 2021). The association of GM130 with RNA (Figure 2) and the potential of GM130 to self-associate and/or phase separate (Rebane et al., 2020) let us to test the capacity of GM130 to assemble RNA-dependent condensates in cells. Transient transfection of untagged full-length GM130 localized to the Golgi (Figure 5 A). After Golgi binding was saturated, excess GM130 entered the nucleus due to an importin α binding site we previously characterized (Wei et al., 2015). At high concentrations, nuclear GM130 assembled into distinct droplet-like structures. The size of the droplets increased with the level of overexpression, and at very high concentrations the droplets tended to blend into an amorphous gel-like structure (Figure 5 A, B). These concentration-dependent condensates of GM130 also formed rapidly within 2 h when expressed by microinjection of the encoding plasmid (Figure 5 C, D).

**Fig.5.**
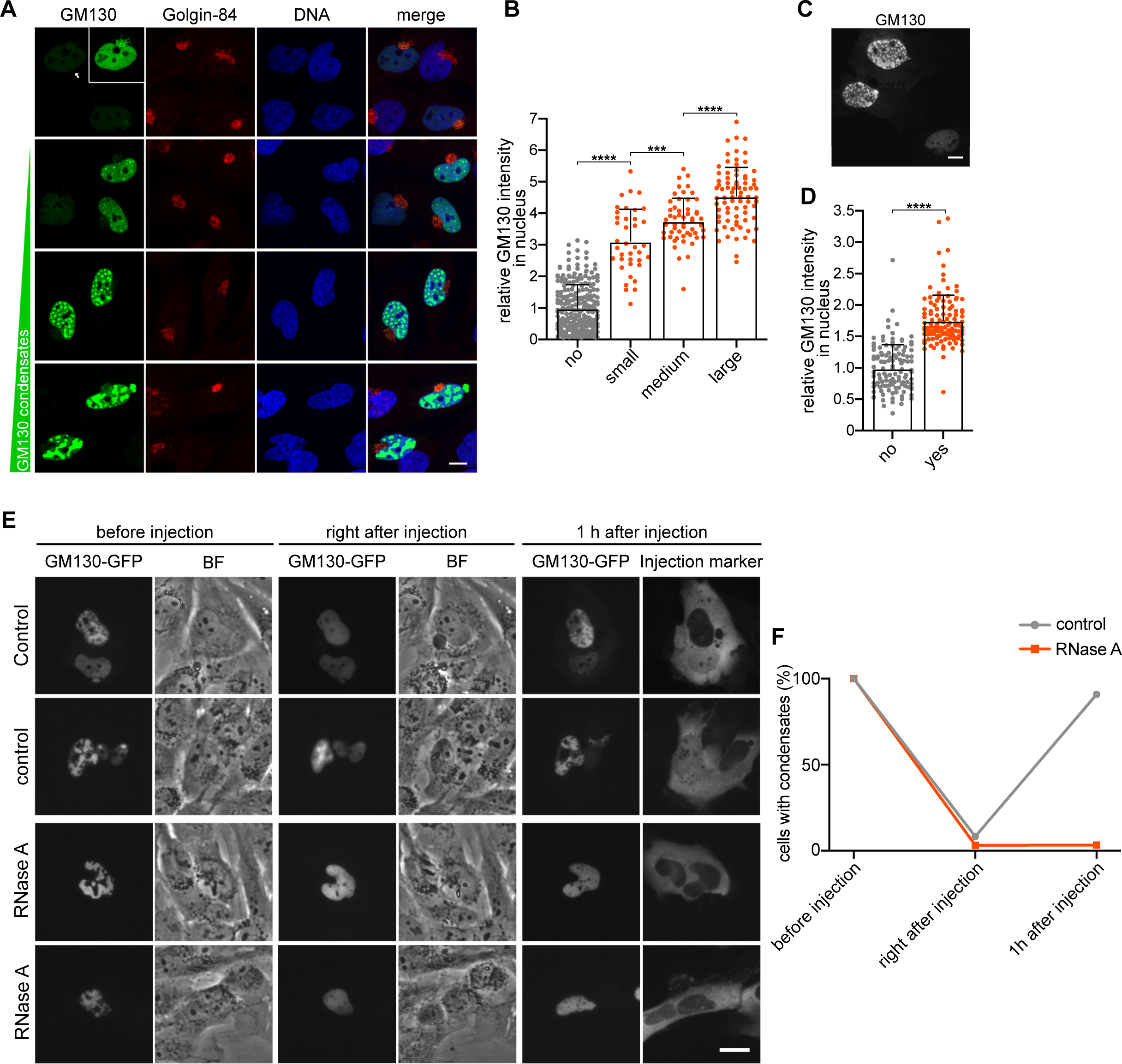
GM130 forms RNA-dependent liquid-like condensates in cells. (A-B). GM130 forms concentration-dependent condensates. Full length GM130 was transiently overexpressed in SV589 overnight and immunolabeled for GM130 (green), Golgi-84 (red) and stained for DNA (blue). Arrow points at the cell with brightness increased in the inset to visualize Golgi-localized GM130. Cells with GM130 condensates were classified into small, medium and large structures. The mean intensity of GM130 in nucleus with or without condensates were quantified in (B). n = 3 independent experiments with the total number of cells analyzed as follows: no condensates: n = 246 cells, small: n = 38 cells, medium: n = 54 cells; large: n = 78 cells. **** P < 0.0001. Error bars represent mean ± SD. Scale bar, 10 µm. (C-D). GM130 forms condensates within 2 h. Full length GM130 was expressed by microinjection of a cDNA encoding plasmid into the nuclei of SV589 cells. Cells were fixed 2 hours later and immunostained for GM130. The average intensity of GM130 in nucleus were quantified as shown in (D). n = 3 independent experiments with the total number of cells analyzed as follows: no: n = 116 cells, yes: n = 114 cells. **** P < 0.0001. Error bars represent mean ± SD. Scale bar, 10 µm. (E). The formation of GM130 condensates in cells depends on RNA. SV589 cells overexpressing GM130-GFP were microinjected with 1 mg/ml RNase A together with fluorescent dextran as an injection marker or fluorescent dextran only as control. Phase contrast images (BF) and images of GFP fluorescence were captured immediately after the injection. The same cells were imaged again after 1 h. Note that condensates were dissolved after injection and reformed within 1 h in control cells but not in RNase A injected cells. Scale bar, 20 µm. (F). Quantitation of the percentage of cells with condensates immediately after injection and after 1 hour from (E). n = 3 independent experiments with the total number of cells analyzed as follows: control: n = 33 cells, RNase A: n = 30 cells.

We then analyzed the dynamics of GM130 condensates by FRAP of GM130 tagged with GFP. Within 20 seconds after photobleaching more than 70% of the GFP signal was restored, indicating liquid-like properties of the condensates (Figure S5 A, B). Furthermore, GM130-GFP droplets were dynamic and frequently merged into larger structures (Figure S5 C).

Next, we tested whether GM130 droplets are RNA-dependent liquid-like condensates. We transiently expressed GM130-GFP and microinjected the cells with buffer or RNase A. GM130 droplets dispersed instantly upon injection of either buffer or RNase A (Figure 5 E, F). However, while droplets in control cells reformed within 60 min, GM130-GFP in RNase injected cells did not reform droplets but remained dispersed. These results show that GM130 has the capacity to assemble RNA-dependent liquid-like condensates in cells that dynamically exchange molecules with the surrounding.

### The N-terminal domain of GM130 is intrinsically disordered and undergoes liquid-liquid phase separation

The extensive coiled-coil domains of GM130 are preceded by an N-terminal domain of 122 residues (N122) predicted to be an intrinsically disordered region (IDR) by IUPred3 (Erdős et al., 2021) (Figure 6 B). IDR-containing proteins play a key role in LLPS and are frequently found in association with RNA in phase separated condensates such as stress granules (Banani et al., 2017; Uversky, 2021). While purifying His-tagged recombinant N122 GM130, we observed that at high concentrations the protein solution turned turbid or opalescent after lowering the buffer salt concentration to 150 mM. Light microscopy revealed that the protein had phase-separated from the bulk solution into distinct spherical droplets (Figure 6 A). As shown in Figure 6D, droplets of monomeric N122-His were only observed at concentrations exceeding the estimated endogenous GM130 concentration of 186 µM on the Golgi membrane in cells (Materials and Methods). We reasoned that LPPS of N122 is diminished by the lack of the coiled coil domains of GM130 that mediate its self-oligomerization into a parallel dimer or tetramer (Ishida et al., 2015). To preserve the valency, we fused N122 to short coiled-coil sequences that assemble into parallel homo-dimers or -tetramers, respectively (Khairil Anuar et al., 2019) (Figure 6 C). We verified the oligomeric state N122 (monomer), N122-Di (dimer) and N122-Tet (tetramer) by chemical crosslinking (Figure S3 A) and then examined the phase separating behavior by DIC microscopy in physiological buffer without any molecular crowding agents. The protein concentration of dimers and tetramers at which droplets formed was substantially lower compared to monomers (Figure 6 D, E). While dimers started to form droplets at 100 µM and tetramers at 12.5 µM, monomers did not phase separate at 100 µM.

**Fig. 6.**
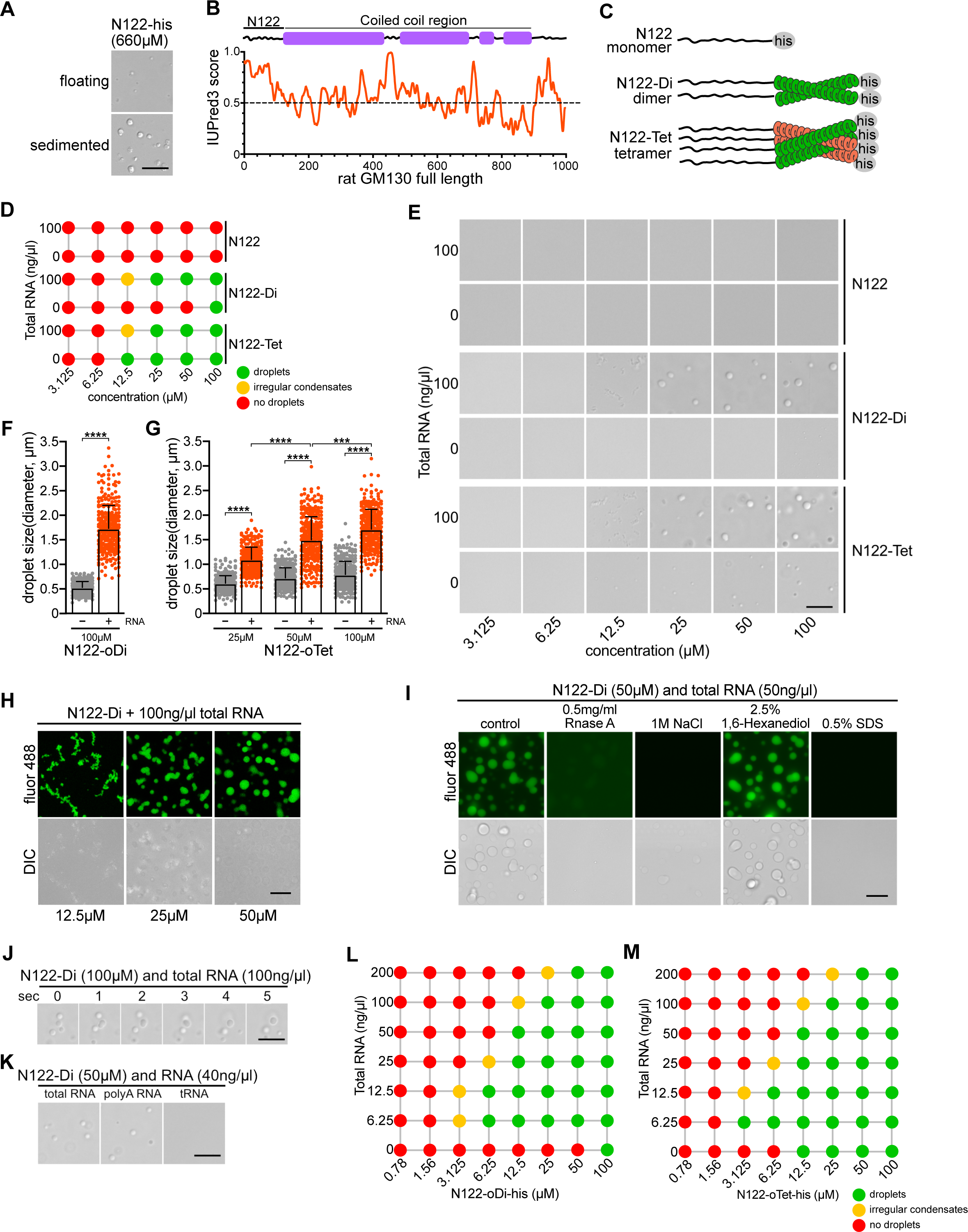
**The N-terminal domain of GM130 forms RNA-dependent condensates *in vitro*** (A). The N-terminal domain of GM130 (N122) forms condensates at high concentration. Recombinant N122-his undergoes liquid-liquid phase separation (LLPS) at high concentration (660 µM). Floating condensates were imaged by DIC microscopy within 5 min after LLPS induction. Sedimented droplets were imaged after 30 min. Scale bar, 10 µm. (B). The N-terminal domain of GM130 of 122 amino acids is highly disordered. GM130 domain organization is aligned with its disordered information by IUPred3. (C). Schematic illustration of monomeric N122, fused to a dimerization domain (N122-Di) and or linked to a tetramerization domain (N122-Tet). (D-E). RNA promotes LLPS of GM130 N122. LLPS of N122, N122-Di or N122-Tet at different concentrations in the presence or absence of 100 ng/µl total RNA from SV589 cells. DIC images of droplets were captured within 5 min after LLPS induction and classified in (D). Representative images captured within 5 min are shown in (E) except for 12.5 µM N122-Di and N122-Tet with RNA which were captured after 20 min. Scale bar, 10 µm. (F). Quantitation of the size of droplets formed by 100 µM N122-Di with or without RNA from (E). n = 318 (no RNA) and 321 (RNA) droplets from three independent experiments analyzed. **** P < 0.0001. Error bars represent mean ± SD. (G). Quantitation of the droplet size formed by N122-Tet at indicated concentrations with or without RNA from (E). n = 3 independent experiments with >100 droplets analyzed per experiment per condition. *** P < 0.001; **** P < 0.0001. Error bars represent mean ± SD. (H). LLPS is concentration dependent. N122-Di (containing 5% fluor 488-labeled N122-Di) and 100 ng/µl total RNA were incubated in 96 well glass bottom plate and droplets were allowed to sediment on glass for 30 min. Representative fluorescent and DIC images were captured after 30 min. Scale bar, 10 µm. (I). N122 droplets are dissolved by RNase and salt. Droplets of N122-Di (containing 5% fluor 488-labeled N122-Di) and total RNA were allowed to sediment on glass for 30 min and then incubated with the indicated reagents. Representative fluorescent and DIC images were taken in 5min after addition of the indicated reagents. Scale bar, 10 µm. (J). Representative DIC images at indicated time points show the fusion of droplets formed by N122-Di and total RNA from SV589 cells. Scale bar, 10 µm. (K). Poly(A) RNA and total RNA support droplet formation, but not tRNA. DIC images of LLPS of N122-Di and total RNA, poly(A) RNA or tRNA. Scale bar, 10 µm. (L). LLPS of N122-Di at different concentrations with increasing concentrations of total RNA from SV589 cells. (M). LLPS of N122-Tet at different concentrations with increasing concentrations of total RNA from SV589 cells.

### RNA stimulates phase separation of N122 GM130

RNA binding proteins with intrinsically disordered protein domains frequently phase separate in combination with RNA (Roden and Gladfelter, 2021). As shown in Figure 2, RNA is bound to GM130 in cells and the N-terminal domain of GM130 is sufficient to bind RNA *in vitro*. We therefore tested the extent to which N122 droplet formation is affected by total RNA isolated from SV589 cells (Figure 6 D, E). We initially used 100 ng/µl RNA, which is in the concentration range of RNA in the cytoplasm of 234 ng/µl (Maharana et al., 2018). RNA strongly promoted droplet formation, which was suppressed by RNase A showing that RNA enhances N122 GM130 phase separation under physiological conditions (Figure S3 B). RNA further lowered the phase transition concentration of N122 dimers to 12.5 µM, which is below the estimated GM130 concentration of about 186 µM on Golgi cisternae in cells (Materials and Methods). Furthermore, RNA significantly increased the volume of the droplet phase, as measured by an increased droplet diameter (Figure 6 F, G). To confirm that the droplets contain N122 protein, we included fluorescently labelled N122-Di in the reaction (Figure 6 H).

Preformed droplets dispersed rapidly by RNase, 1M NaCl or 0.5% SDS confirming that they are liquid-like condensates (Figure 6 I). Droplets were however not affected by 1,6-hexanediol treatment, a compound that can dissolve condensates considered of weak hydrophobic interactions (Kroschwald et al., 2017). In agreement with condensates in cells (Figure S2), droplets of N122-Di and RNA coalesced into larger spherical droplets, reflecting their liquid-like properties to reduce the surface area in correlation to the volume (Figure 5 J). We next tested if different kinds of RNAs differ in their abilities to stimulate N122 droplet formation. The poly(A) RNA fraction of total RNA induced droplets as potently as total RNA, while tRNA did not promote N122-Di condensation (Figure 6 K).

We observed that increasing amounts of RNA stimulated droplet formation (Figure 6 L, M). The greatest extent of droplet formed at a ratio of N122 to RNA of 25 µM:100 ng/µl, which is within the estimated physiologic concentrations of GM130 on the Golgi and RNA in the cytoplasm. Further increase of RNA concentrations suppressed condensates showing that the protein RNA ratio is of importance. Too low or too high ratios disfavor droplet formation, which is has been reported for various RBPs such as TDP-43, Whi3 and FUS (Burke et al., 2015; Zhang et al., 2015; Maharana et al., 2018). These results show that the intrinsically disordered N-terminal domain of GM130 undergoes RNA-dependent liquid-liquid phase separation under physiological relevant conditions.

### N122 GM130 and RNA are sufficient to link Golgi membranes in vitro

Our results show that acute degradation of GM130 or RNA in cells leads to unlinking of Golgi ribbon stacks (Figure 3). Similarly, oxidative stress displaces RNA from GM130 and causes Golgi fragmentation, both of which are restored after arsenite washout (Figure 4). Furthermore, GM130 undergoes RNA-dependent phase separation in cells (Figure 5) and *in vitro* (Figure 6). Considered together, our results indicate that molecular condensates of GM130 and RNA are joining adjacent Golgi stacks. To determine whether droplets of N122 and RNA are sufficient to link Golgi stacks, we performed the condensation reaction in the presence of Golgi membranes. Purified rat liver Golgi membranes were fluorescently stained with the red lipophilic dye DiD perchlorate, incubated with OG488 labelled N122-Di and total RNA and then observed by fluorescence microscopy. As controls, N122 or RNA alone did not change on the morphology or distribution of the Golgi membranes and condensates of N122 were not observed (Figure S4). When Golgi membranes were incubated with N122 and RNA, condensates formed to a similar extent as without Golgi membranes (Figure 6). These condensates formed adjacent to the Golgi membranes in discrete patches decorating the edges. The multiple patches of Golgi membranes were thereby joint by N122 condensates into a string-like pattern (Figure 7 B, C). These results suggest that condensates of RNA, N122 and Golgi membrane proteins are sufficient to link Golgi stacks into a string resembling ribbon-like structure *in vitro*.

**Fig. 7.**
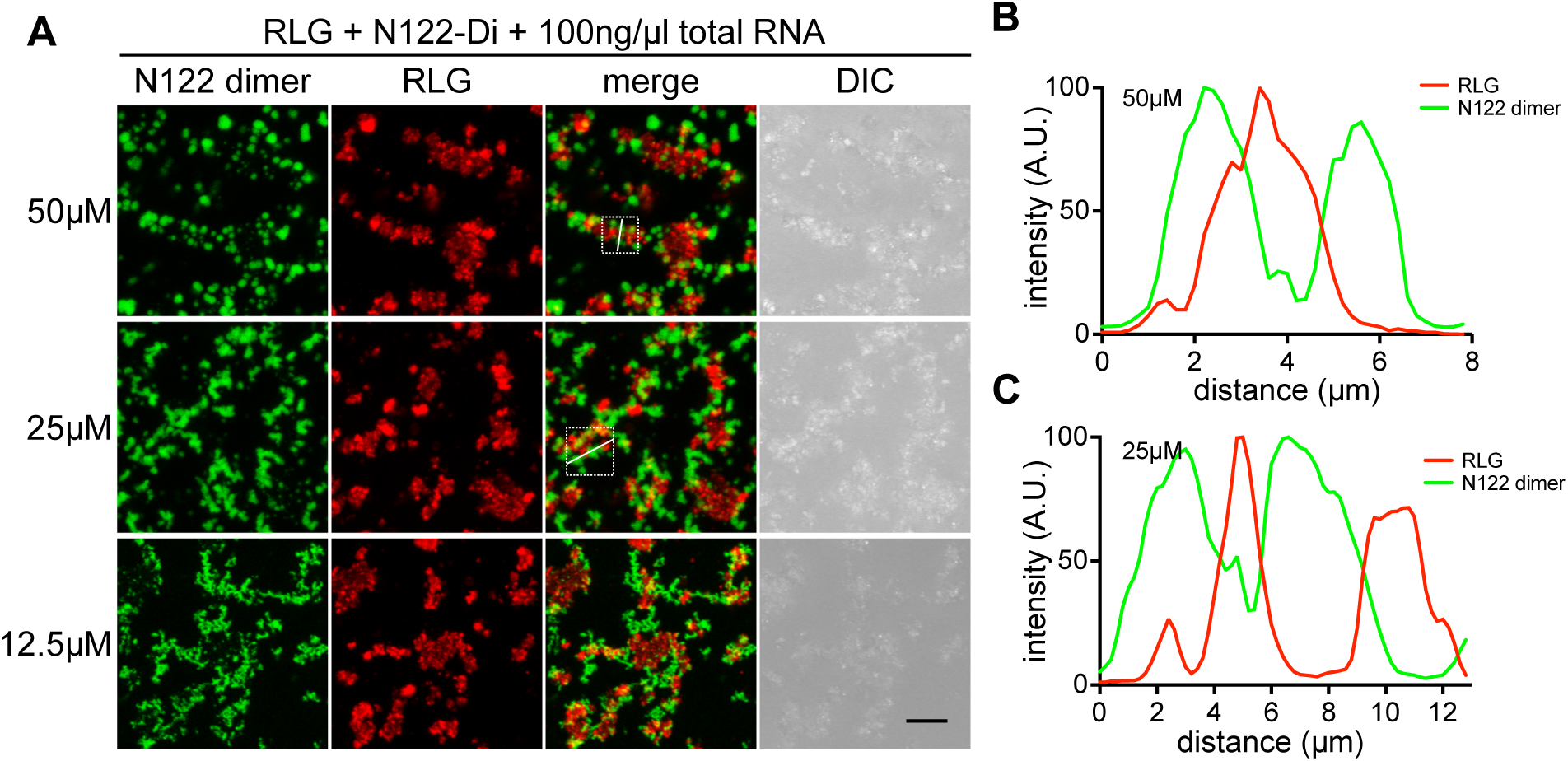
**Condensates formed by N122-Di and RNA are sufficient to link Golgi membranes *in vitro*.** (A-C). Fluorescently stained purified rat liver Golgi membranes (RLG) were incubated with N122-Di (containing 5% fluor 488-labeled N122-Di) and 100 ng/µl total RNA from SV589 cells. Images were captured 30 min after incubation for sedimented RLG. Insets show the white lines marking the line scan of the fluorescence intensities shown in (B) and (C), respectively. Scale bar, 10 µm.

## Discussion

Here, we identify the Golgi resident protein GM130 as a novel membrane-bound RNA binding protein. GM130 contains an intrinsically disordered domain at its N-terminus (N122), which is sufficient to recruit RNA to drive droplet-like biomolecular condensate assembly *in vitro*. Overexpressed of GM130 in cells further support our *in vitro* data that GM130 assembles into RNA-dependent liquid-like condensates. RNA addition lowered the threshold of GM130 phase separation while RNA loss caused by RNase digestion disrupted GM130 condensation both *in vitro* and in cells. Our key functional experiment, namely laterally linking of stacks, is a reconstruction of aspects of lateral linking of stacks into a ribbon-like structure. Similarly, acute loss of either RNA or GM130 leads to a disjointed Golgi ribbon in cells. Under stress conditions, RNA and associated RBPs are lost from GM130, and the ribbon is disconnected. When cells returned to normal growth conditions RNA binding to GM130 and ribbon linking is fully restored. Thus, these results indicate that the Golgi ribbon is tied together by a functional condensate of GM130 and RNA. As such, RNA polymers can be considered as structural elements that scaffold the Golgi by associating with membrane anchored GM130.

GM130 was previously identified to associate with RNA in an unbiased screen aimed to identify the human RNA binding proteome, but direct RNA binding was not further verified (Trendel et al., 2019). GM130 further shares characteristic sequence motifs that are found in RBPs that have tendencies to form condensates: a RNA binding site, a weakly self-associating intrinsically disordered protein region (IDR), and an oligomerization domain (Sanders et al., 2020). The N-terminal 122 amino acids of GM130 are predicted to form an IDR and has the tendency to self-associate into droplets, although only at levels that exceed the cellular concentrations. However, the threshold of N122 droplet formation is greatly lower by RNA binding. A previous study describing the potential of GM130 to phase separate did not report RNA as an essential driver and the functional significance was undefined (Rebane et al., 2020). It is however likely that the protein already had RNA bound as it was purified from tissue culture cells.

GM130 forms dimers and tetramers on the surface of the Golgi membrane (Ishida et al., 2015). The valency mediated by the extensive coiled-coil domains of GM130 greatly increases its tendency to undergo RNA dependent LLPS (Figure 6 C-E). This is a common feature of RBPs that are found in phase separated structures. The multi valency of the protein allows for the cross linking of long RNA molecules which stimulates phase separation. This has been exemplified by stress granule nucleation factor G3BP1. Its dimerization and RNA binding are required for phase separation into stress granules (Sanders et al., 2020), while deletion of the G3BP dimerization domain suppresses granule formation. Furthermore, stress granules function as part of the antiviral response and various viruses cleave or inhibit G3BP1 dimers to prevent granule assembly allowing the virus to escape the innate immune response (Corbet et al., 2022).

In addition to valency, restricting GM130 localization to the 2D membrane surface of the Golgi increases the local concentration thereby favoring its ability to phase separate. Membrane confinement of GM130 raises its local concentration by more than one order of magnitude from 146 nM to 186 µM (see Methods). This concentration is within the range we found to be effective for LLPS of N122 and RNA *in vitro*. We further observed most effective LLPS of N122 at concentrations that are within the range of cytoplasmic RNA levels (Maharana et al., 2018).

Attaching proteins to the membrane not only facilitates phase separation due to an increase in local concentration, it also limits condensate size by restricting protein mobility (Snead et al., 2022). While droplet formation reconstituted *in vitro* assemble into micron sized structures, condensates in cells are usually of much smaller size of less than 100-300 nm in diameter (Lyon et al., 2021). On the surface of the Golgi, these size-restricted islands of GM130 condensates may be interconnected by long mRNA molecules into a grid-like meshwork that is anchored on the membrane surface by GM130. The propensity to form condensates might not be restricted to GM130. Other members of the Golgin protein family of Golgi-associated coiled-coil proteins have the potential to form droplets when overexpressed in cells (Ziltener et al., 2020). Whether these Golgins also recruit RNA needs to be determined, but RNA binding may not only link islands of homotypic condensates on the same cisternae, but also between cis, medial and trans cisternae. The RNA-linked islands of Golgin condensates then collectively forms a meshwork or matrix forming a polymeric exoskeleton surrounding the Golgi apparatus. In fact, members of the Golgin and GRASP protein families jointly act as a matrix that stabilized Golgi membranes through multimeric interactions and defines the higher order structure of the Golgi ribbon (Slusarewicz et al., 1994; Seemann et al., 2000). We recently showed that acute depletion of either GRASP65 or GRASP55 does not affect the Golgi structure, while removal of both GRASPs indirectly leads to the slow gradual loss of several Golgins. This showed that the multi valent matrix meshwork is robust enough to compensate for the loss of one component, while elimination of two component sufficiently weakens the collective interactions of the network to gradually destabilize of the Golgi ribbon integrity (Zhang and Seemann, 2021). Our current data suggests that RNA may be an integral component of the matrix to link the Golgi matrix proteins together into a meshwork in a process that is driven by condensate formation of Golgins.

We found that GM130 directly recruits poly(A) RNA, which raises the question whether the transcripts are locally translated at the Golgi. Associated with the RNA component of the complex are RBPs including G3BP1 and FXR1. These RBPs are found in stress granules where they are thought to incorporate non-translated mRNAs thereby suppressing translation (Khong et al., 2017). Although these RBPs are predominantly cytosolic, they are enriched on purified Golgi membranes and can be found in proximity to GM130 on the Golgi in cells (Figure 1). Furthermore, we detected some FXR1 that tends to accumulate in discrete foci in the prenuclear Golgi region close to the GM130 signal in non-stressed cells. Similar to stress granules, the clustering of FXR1 in foci associated with the Golgi suggests that ribosome recruitment and local translation of the RNA is suppressed.

On the other hand, EM studies described polyribosomes in the Golgi area of rat hepatocytes and on enriched rat liver Golgi membranes. However, unlike ribosomes on ER membranes, these polysomes are only close to membranes and not directly membrane attached (Elder and Morré, 1976). Other EM studies showed that Golgi membranes are surrounded by proteinaceous structures protruding from the cisternal membrane bilayer (Cluett and Brown, 1992). This matrix defines a "zone of exclusion" that is scarce or devoid of organelles and ribosomes (Mollenhauer and Morré, 1978; Farquhar and Palade, 1981; Morré and Mollenhauer, 2008). The lack of Golgi membrane-bound ribosomes and the association of granule-forming RBPs with GM130 thus argues that GM130-bound RNA is not translated. Whether transcripts bound to GM130 and/or other Golgi resident proteins are translationally active or inactive requires further experimental investigation. We propose that GM130 differs from other membrane-associated RBPs that capture RNA for local organelle-coupled translation (Béthune et al., 2019). Instead, RNA recruited to GM130 is not used for protein synthesis, but functions as a molecular scaffold that organizes the underlying membrane by favoring condensate assemblies with GM130 to link Golgi membranes into a ribbon. The expression and spatio-temporal distribution of the RNA species, such as long non-coding RNA or mRNA of different lengths and structures, further has the potential to dynamically modulate the extent, structure and surface of the condensate and thereby their function (Ma et al., 2021).

Several lines of evidence showed that GM130-RNA condensates function to dynamically link stacks into a ribbon. Acute degradation of either RNA or GM130 disrupts ribbon integrity. Similarly, exposure of cells to oxidative stress conditions displaces RNA from GM130 and disconnects the ribbon, which is both fully restored after the cells returned to normal growth conditions. Importantly, we have devised a defined *in vitro* assay using purified Golgi membranes to reconstitute the lateral linking of stacks by condensate formation of N122 GM130 and RNA. Strikingly, the membranes are not engulfed by condensates, but droplets formed in discrete patches decorating the edges of the Golgi membranes. The locally attached droplets bridge adjacent membrane patches into a string-like pattern resembling a ribbon.

Cell-free reconstitution systems are powerful approaches in determining molecular mechanisms of cellular function. Several aspects of Golgi organization have been reconstituted *in vitro*, including vesicle budding and fusion, mitotic unstacking and vesiculation and post-mitotic reformation of cisternae into stacks. However, because RNA was not known to be the key factor, lateral linking of stacks could so far not be reconstituted, and studies aimed to determine the molecules that link stacks into a ribbon in cell mainly relied on genetic perturbations including RNAi knockdown or gene deletions. However, long-term chronic perturbations often lead to indirect phenotypes. Given that the Golgi is a dynamic structure that is constantly remodeled by budding and fusion of transport vesicles, chronic perturbation of Golgi proteins is often associated with indirect morphological changes. Downregulation of GRASP55 was reported to unlink the Golgi ribbon (Feinstein and Linstedt, 2008; Xiang and Wang, 2010), but we previously reported that acute elimination of GRASP55 could not phenocopy ribbon unlinking, and ribbon integrity was only compromised after long-term depletion (Zhang and Seemann, 2021). Similarly, RNAi of GM130 let to the loss of GRASP65, which in turn resulted in ribbon fragmentation associated with shortened cisternae and with accumulated vesicles around the stacks (Puthenveedu et al., 2006). To circumvent pleiotropic effects, we acutely degraded endogenous GM130 by Trim-away, eliminated RNA by RNase in cells, and reconstituted linking of Golgi membranes *in vitro*. Together, these lines of evidence support the model that RNA is a structural element that forms meshwork with GM130 to maintaining the integrity of the Golgi ribbon.

## Materials and Methods

### Cell culture and drug treatments

HEK293T (ATCC), NRK (normal rat kidney) (ATCC), SV589 (SV40 immortalized human fibroblasts) (Yamamoto et al., 1984), RC55 and RC65 cells were cultured at 37°C and 5% CO_2_ in medium A (DMEM (Corning), 10% cosmic calf serum (HyClone), 100 units/ml penicillin (GoldBio) and 100 µg/ml streptomycin (GoldBio)). For microinjection experiments, the medium was switched to medium B (medium A supplemented with 50 mM HEPES-KOH pH 7.4).

RC55 and RC65 are SV589 cell lines stably expressing doxycycline-inducible OsTIR1-2xMyc as well as endogenous GRASP55 or GRASP65 tagged with a Flag epitope followed by a minimal auxin inducible degron sequence (3xFlag-mAID) (Zhang and Seemann, 2021). Expression of the E3 ubiquitin ligase component OsTIR1-2xMyc was induced by 0.5 μg/ml doxycycline (Sigma) for 6 h. GRASP65 was then depleted by treatment with 0.5 mM of the auxin indole-3-acetic acid (IAA, Abcam) for 2 h (Zhang and Seemann, 2021).

NRK/TRIM21 cells stably expressing mCherry-TRIM21 were generated by transducing NRK cells with lentiviral particles that were produced by co-transfection of HEK293T cells with pSMPP- mCherry-hTRIM21 (a gift from Leo James, Addgene # 104972), psPAX and pVSVG using PEI Max (Polysciences). Single clones were isolated by limited dilution and expression of mCherry- TRIM21 was verified by fluorescence microscopy.

To generate SV589/NAGTI-GFP and NRK/TRIM21/NAGTI-GFP cell lines that express the Golgi marker NAGTI-GFP, we transduced SV589 or NRK/TRIM21 cells with lentivirus particles generated in HEK293T by co-transfection with pLVX-NAGTI-GFP-Puro (Zhang and Seemann, 2021), psPAX and pVSVG. NAGTI-GFP expressing cells from a mixed population were used for FRAP experiments.

Stress granules were induced by treating SV589 cells with 250 µM sodium arsenite (Sigma) for 1 h.

GM130 immunoprecipitates from SV589 cell lysate or N74-strep pulldowns from cell lysate were treated with 5-50 µg/ml RNase A (Sigma). The RNase inhibitor RNasin Plus (Promega) was used at 40 U/ml.

### Plasmids

All oligonucleotide sequences are listed below. To construct pET23a(+) GM130 N74-strep, the GST sequence in pET23a(+) GM130 N74-GST (Wei et al., 2015) was replaced with a sequence encoding Strep-tag II (WSHPQFEK) derived by annealing of oligos #473 and #474.

Full-length rat GM130 (NP_072118.2) with the C-terminus fused to GFP was generated by amplifying GM130 by PCR from pcDNA3.1(-) GM130 (Wei et al., 2015) using primers #509/#24 and cloning into pcDNA3.1(-) to obtain pcDNA GM130 FL. Full length GM130 was then amplified by PCR with primers #510/#508 and ligated into pEGFP-N1 to yield pEGFP-GM130 FL.

pET23a(+) GM130 N122-his was produced by replacing the N74 sequence of pET23a(+) GM130 N74-his (Wei et al., 2015) with the sequence encoding the N-terminal 122 amino acids of GM130 amplified from pcDNA GM130 FL by PCR using primers #500/511. The oDi sequence encoding a coiled-coil dimerization domain was amplified by PCR (#503/#504) from SpyCatcher002-oDi (a gift from Mark Howarth, Addgene # 124661) (Khairil Anuar et al., 2019) and then cloned into pET23a(+) GM130 N122-his to yield pET23a(+) GM130 N122-oDi-his. Similarly, the oTet sequence encoding a tetramerization coiled-coil domain was obtained by PCR (#505/#506) from SpyCatcher002-oTet (a gift from Mark Howarth, Addgene # 124663) (Khairil Anuar et al., 2019) and cloned into pET23a(+) GM130 N122-his to construct pET23a(+) GM130 N122-oTet-his. The proteins encoded by the sequences of N122-his, N122-oDi-his and N122-oTet-his are referred to as N122, N122-Di and N122-Tet, respectively.

### Protein purification

GM130 N74-strep was expressed in BL21-Gold(DE3)pLysS *E. coli* (Aglient). GM130 N122, N122- Di and N122-Tet were expressed in Rosetta 2(DE3) *E. coli* (MilliporeSigma). Protein expression was induced with 0.5 mM IPTG for 3 h at 37°C except N122-Tet which was induced overnight at 18°C. Cells were pelleted, resuspended in HSB (PBS supplemented with 150 mM NaCl, 1 mM DTT, protease inhibitor cocktail Complete (Roche)), lysed with an EmulsiFlex-C5 homogenizer (Avestin) and lysates were cleared by centrifugation. N74-strep was bound on Strep-TactinXT 4Flow beads (IBA), washed with strep buffer (100 mM Tris-HCl pH 8.0, 150 mM NaCl, 1 mM EDTA), and eluted with 50 mM biotin (Santa Cruz) in strep buffer. His-tagged proteins were bound by Ni-NTA agarose (Qiagen) and eluted with 500 mM imidazole in HSB. Eluted proteins were desalted on a 10DG column (Bio-Rad) into 50 mM HEPES-KOH pH 7.4, 375 mM NaCl, 1mM DTT, concentrated with a 3 kDa Amicon Ultra centrifugal filter (Millipore), flash frozen in liquid nitrogen and stored at -80°C.

### Immunoprecipitation and mass spectrometry

SV589, RC55 or RC65 cells were lysed for 30 min on ice in lysis buffer (50 mM Tris-HCl pH 7.4, 150 mM NaCl, 1 mM EDTA, 1% NP40, protease inhibitor cocktail Complete (Roche)) and lysates were cleared by centrifugation for 10 min at 16,000 g at 4°C. Flag-tagged proteins were bound to anti-FLAG M2 affinity gel (Sigma-Aldrich) for 1 h at 4°C. Beads were washed with lysis buffer and Flag-tagged GRASP55 and GRASP65 were eluted with 0.5 mg/ml 3xFLAG peptide (Apexbio) in lysis buffer. Eluted proteins were separated by SDS-PAGE, cut out protein bands were digested with trypsin and processed by a short reverse-phase LC-MS/MS on a Orbitrap Fusion Lumos Mass Spectrometer (Thermo Fisher Scientific). Proteins were identified using Proteome Discoverer 2.4 software (Thermo Fisher Scientific).

Endogenous GM130 was immunoprecipitated by incubation of cleared SV589 cell lysates for 1 h at 4°C with either 1 µg rabbit anti-GM130 antibodies (Proteintech), 5 µl rabbit polyclonal anti- GM130 serum (Wei and Seemann, 2009b) or 1 µg rabbit IgG (ImmunoReagents) as control. Complexes were bound on 10 µl Dynabeads Protein A (Invitrogen) or Protein A/G Plus agarose (Santa Cruz) for 1 h at 4°C. Beads were washed with lysis buffer and either boiled in SDS sample buffer for Western blotting, treated for 30 min at 4°C with 20 µg/ml RNase A (Sigma) or 40 U/ml RNase inhibitor (RNasin Plus; Promega) in lysis buffer followed by Western blotting, or beads were subjected to RNA extraction using a Direct-zol RNA miniprep kit (Zymo Research).

### Strep-tag pulldown assay

SV589 cells were lysed in lysis buffer for 30 min on ice. Cleared lysates were mixed with 7 µg N74-strep for 1 h at 4°C and then incubated with 10 µl slurry Strep-TactinXT 4Flow beads (IBA) for 1 h at 4°C. Washed beads were treated with RNase A in lysis buffer for 30 min at 4°C, washed, and proteins were eluted by boiling in SDS sample buffer.

For *in vitro* binding of N74-strep and G3BP1-his in the presence of RNA, 4.2 µg N74-strep and 10 µg G3BP1-his were mixed with total RNA isolated from SV589 cells in 20 mM HEPES-KOH pH 7.4, 150 mM NaCl, 5% glycerol, 1 mM DTT, 40 U/ml RNasin Plus. The reaction was incubated for 20 min at room temperature followed by 10 min on ice and then incubated with 10 µl slurry Strep-TactinXT 4Flow beads for 30 min at 4°C. The beads were washed with 20 mM HEPES-KOH pH 7.4, 150 mM NaCl, 5% glycerol, 1 mM DTT, 10 U/ml RNasin Plus and bound proteins were eluted by boiling in SDS sample buffer.

### Microinjection

Cells grown on glass coverslips or 35 mm glass bottom dishes (MatTek Corporation) were microinjected using a FemtoJet microinjector (Eppendorf) and a micromanipulator 5171 (Eppendorf). The medium was exchanged to medium B before injections. Following the injections, cells were switched back to medium A and incubated at 37°C and 5% CO_2_.

Acute degradation of endogenous GM130 in NRK/TRIM21/NAGTI-GFP cells by Trim-Away (Clift et al., 2017) was induced by microinjection of 0.5 mg/ml affinity purified rabbit anti-GM130 antibodies (morsel) (Wei et al., 2015) or 0.5 mg/ml anti-DDDDK IgG (Bio X Cell) as control mixed with 2 mg/ml Texas-Red dextran (Invitrogen) in KH buffer (20 mM HEPES-KOH pH 7.4, 50 mM KCH3COO). Injected cells were incubated for 2 h and either subjected to FRAP analysis or fixed in 3.7% formaldehyde in PBS for immunofluorescence analysis.

SV589 cells were microinjected with 1 mg/ml RNase A (Sigma) mixed with 2 mg/ml fluorescein dextran (Invitrogen) in 25 mM HEPES-KOH pH 7.4. Injected cells were incubated for 1 h and fixed in 3.7% formaldehyde in PBS for immunofluorescence analysis.

To determine RNA dependency of GM130 condensates in cells, SV589 cells grown on glass coverslips were transfected with pEGFP-GM130 FL using PEI Max (Polysciences) and incubated overnight. Cells with GM130-GFP condensates were imaged and then injected with 1 mg/ml RNase A together with 2 mg/ml Texas-Red dextran or 2 mg/ml Texas-Red dextran in 25 mM HEPES-KOH pH 7.4. Injected cells were imaged immediately after injection, incubated on the microscope stage for 1 h at 37°C and imaged again. Phase-contrast pictures were captured with a LD-A-PLAN 20x/0.3 Ph1 objective (Zeiss) and GFP pictures were acquired with a Plan-Apochromat 20x/0.6 objective (Zeiss).

### Immunofluorescence and Microscopy

Cells grown on glass coverslips were fixed for 12 min at RT in 3.7% formaldehyde in PBS and permeabilized for 15 min in -20°C methanol. Cells were washed with PBS and incubated with indicated primary antibodies for 30 min at 37°C followed by Alexa Fluor-conjugated secondary antibodies for 30 min at 37°C. DNA was stained with 1 µg/ml Hoechst 33342 (Invitrogen) in PBS for 5 min at RT and cells were mounted in Mowiol 4-88 (Calbiochem) solution (Wei and Seemann, 2009a). Images shown in Fig.1H, 1K,1M, 3D, S1A, 4A, 5A were acquired using a LSM880 inverted confocal microscope (Zeiss) with a Plan-Apochromat 63x/1.4 objective (Zeiss) or 40x objective (Zeiss). Z-sections were captured at 0.47 µm (63x objective) or 1 µm (40x objective) intervals. Maximum intensity projections of z-stacks are presented in the figures. Images in Fig. 3A, 3H, 3J, 5C were captured with an Axiovert 200M microscope (Zeiss), an Orca 285 camera (Hamamatsu), Openlab 4.0.2 software (Improvision) and a Plan-Neofluar 40x/1.3 DIC objective (Zeiss).

### Proximity Ligation Assay (PLA)

SV589/NAGTI-GFP cells grown on glass coverslips were transfected with siRNA Universal Negative Control #1 (Sigma) or duplexes targeting GM130 or FXR1 (Sigma) using Lipofectamine RNAiMAX (Invitrogen). Cells were fixed 72 h after transfection in -20°C methanol and subjected to PLA using the Duolink In Situ Red Starter Kit Mouse/Rabbit (Sigma) according to the manufacturer’s instructions. Briefly, fixed cells were blocked, incubated with mouse monoclonal IgG against FXR1 (Sigma) and affinity purified rabbit anti-GM130 (morsel) for 30 min at 37°C. Cells were then washed, incubated with anti-mouse MINUS and anti-rabbit PLUS PLA probes for 1 h at 37°C followed by ligation and amplification to generate a Texas Red PLA fluorescence signal. DNA was labeled with 1 µg/ml Hoechst 33342 in PBS for 5 min at RT and cells were embedded in Mowiol 4-88 solution. Confocal z-stacks were captured at 1 µm intervals and are shown as maximum intensity projections. The number of PLA puncta associated with the Golgi marked by NAGTI-GFP were counted per cell per condition and normalized to control siRNA. The mean of the control siRNA was normalized to 1.

### Fluorescence recovery after photobleaching (FRAP)

FRAP was performed on NRK/TRIM21/NAGTI-GFP cells that were depleted of GM130 for 2 h by Trim-Away as described under microinjections. SV589/NAGTI-GFP cells were treated with 250 µM Sodium arsenite (SA) for 1 h. SA was washed out and the cells were subjected to FRAP analysis or incubated for 16 h before FRAP analysis. FRAP was performed at 37°C and 5% CO_2_ using a Zeiss LSM880 inverted microscope and a Plan-Apochromat 63x/1.4 objective (Zeiss). The Golgi area as marked by NAGTI-GFP was magnified six times and time-lapse images were acquired at 3 s intervals. The mean GFP fluorescence intensity of the photobleached area and the adjacent area at each time point were measured using Fiji software. The recovery rate was calculated at each time point as the ratio of the mean intensity of the photobleached area to the adjacent area and then normalized to pre-bleach conditions.

For FRAP analysis of GM130 condensates, SV589 cells grown on 35 mm glass bottom dishes were transfected with pEGFP-GM130 FL and incubated overnight. FRAP analysis was performed as above. Cells with GM130 condensates were magnified four times and time-lapse images were acquired each second.

### Crosslinking of recombinant proteins

GM130 N122 proteins were crosslinked using disuccinimidyl suberate (DSS) according to the manufacturer’s instructions (Thermo Fisher). Briefly, 1 mg/ml or 2 mg/ml of N122, N122-Di and N122-Tet protein were incubated with 10 fold molar excess of cross-linker DSS in a final volume of 20 µl for 30 min at RT in 50 mM HEPES-KOH pH 7.4, 375 mM NaCl, 1mM DTT. The reaction was stopped by adding 50 mM Tris HCl pH 7.5 for a minimum of 15 min and analyzed by SDS-PAGE and Coomassie blue staining.

### Isolation of RNA

Total RNA was purified from SV589 cells using the Direct-zol RNA miniprep kit (Zymo Research). poly(A) RNA was isolated from the total RNA with Dynabeads Oligo (dT)_25_ (Thermo Fisher) according to the manufacturer’s instructions. Briefly, 75 µg of total RNA was incubated for 2 min at 65°C and incubated with 1 mg beads for 5 min at RT. Beads were washed, and bound poly(A) RNA was released from the beads by increasing the temperature to 80°C for 2 min.

### Liquid-liquid Phase Separation (LLPS)

LLPS was performed in regular salt buffer (20 mM HEPES, 150 mM NaCl, 1 mM DTT) at RT. GM130 N122, N122-Di or N122-Tet stored in 50 mM HEPES-KOH pH 7.4, 375 mM NaCl, 1 mM DTT were serial diluted in the same buffer and then mixed with 1.5 folds volume of DEPC-treated H_2_O or RNA in DEPC-treated H_2_O to induce LLPS. The samples were mixed in low binding tubes and transferred to a sandwiched chamber composed of a glass coverslip, a glass slide and a double-sided spacer (Yang et al., 2020). Images of floating droplets were captured within 5 min (Fig. 6A, E, J, K and S3B). Sedimented condensates in Fig. 6A were imaged after 30 min. Irregular condensates formed by incubation of 12.5 µM N122-Di or N122-Tet with 100 ng/µl total RNA were imaged after 20 min (Fig. 6E). DIC images were captured with an Axiovert 200M microscope (Zeiss) and a Plan-Neofluar 40x/1.3 DIC objective (Zeiss).

For LLPS of fluorescent proteins, N122-Di was incubated with HIS Lite OG488-Tris NTA-Ni Complex (AAT Bioquest) at 2:1 molar ratio for 30 min on ice. Unbound dye was removed from N122-Di-OG488 by desalting on a 10DG column (Bio-Rad) into 50 mM HEPES-KOH pH 7.4, 375 mM NaCl, 1mM DTT. LLPS was assessed by incubation of N122-Di containing 5% N122-Di- OG488 with 100 µg/ml total RNA and 40 U/ml RNase inhibitor in regular salt buffer. Samples were transferred into a 96 well glass bottom plate (Greiner Bio-One) and incubated for 30 min at room temperature. 7-10 confocal z-sections were captured at 1 µm intervals with a 40x objective and shown as maximum intensity projection (Fig. 6H). LLPS shown in Fig. 6I was performed by incubation of 50 µM N122-Di containing 5% N122-Di-OG488 and 50 µg/ml total RNA in a 96 well glass bottom plate for 30 min before addition of 0.5 mg/ml RNase A, 1 M NaCl, 2.5% 1,6- Hexandiol (Sigma) or 0.5% SDS. Samples were then imaged within 5 min with an Axiovert 200M microscope (Zeiss) and a Plan-Neofluar 40x/1.3 DIC objective (Zeiss).

LLPS in presence of rat liver Golgi (RLG) membranes (Fig.7 and S4) was observed using a LSM880 inverted microscope (Zeiss) with a 40x objective (Zeiss). Rat liver Golgi membranes were isolated as previously described (Tang et al., 2010). Golgi membranes (1.5-2 mg/ml protein) in KHM buffer (20 mM HEPES-KOH pH 7.4, 0.2 M sucrose, 60 mM KCl, 5 mM MgCl_2_, 1 mM DTT, protease inhibitor, 40 U/ml RNase inhibitor) were fluorescently labeled with 2 µM of the lipophilic dye DiD Perchlorate (AAT Bioquest) for 30 min on ice. Labelled Golgi membranes were then mixed with N122-Di and total RNA. Samples were transferred to a 96 well glass bottom plate, incubated for 30 min at room temperature, and imaged by confocal microscopy using a 40x objective at 1 µm z-intervals with 14-16 sections in total. Maximum intensity projections are presented in the figures.

### Image Analysis and Quantitation

Image analysis was performed using Fiji (ImageJ2). Statistical analyses were conducted using Prism 8 or 9 software (GraphPad). The statistical significance was analyzed by Student’s t tests.

The Golgi area in Fig. 3B, 3K and 4B was defined by the fluorescent signal distribution of GM130, GRASP55 or GM130, respectively. The signal was encircled using the Freehand tool and analyzed by the Measure function (Fiji). The Golgi elements per cell in Fig. 3C and 4C labeled by GM130 were selected by applying a fixed threshold and the objects were then counted using the Analyze Particles function (Fiji).

Stacks in Fig. 3D and S3A were defined as Golgin-97 elements that associate with GM130 structures, or TGN38 structures that associate with Golgin-84 by more than 50% of their length (Rabouille et al., 1995). The length of at least ten of the largest Golgin-97 elements per cell and the linking GM130 structures were measured using Fiji and the length of TGN38 and associated Glogin-84 were measured in five random square areas of 4 µm^2^. At least 10 cells per condition were analyzed in each of three independent experiments.

The estimated concentration of endogenous GM130 on the cis Golgi cisternae of 186 µM was calculated by dividing the number of GM130 molecules per cell by the occupied volume on the surface of the cisternae. The number of GM130 molecules per HeLa cell was reported by quantitative mass spec as 86304 (Kulak et al., 2014), which corresponds to n_GM130/cell_ = 1.43 x 10^- 19^ mol. GM130 localization is restricted to the surface of one cis cisternae within a stack (Trucco et al., 2004) with a reported diameter of a cisternae of 0.7 µm (Weidman et al., 1993; Trucco et al., 2004; Koreishi et al., 2013), resulting in a surface area of A = π x r^2^ = 0.385 µm^2^ x 2 surfaces per cisternae = 0.77 µm^2^. The thickness of proteinaceous structures extending from the cisternal membrane bilayer is 11 nm (Cluett and Brown, 1992), which results in an estimated volume of GM130 covering the surface of all Golgi cisternae per cell V_GM130/stack_ = A x 11 nm = 0.77 x 10^-17^ l, multiplied by the average copy number of stacks per cell of 100 (Trucco et al., 2004), V_GM130/cell_ = 0.77 x 10^-15^ l. The molar concentration of GM130 is therefore: c_GM130_ = n_GM130/cell_ / V_GM130/cell_ = (1.43 x 10^-19^ mol) / (0.77 x 10^-15^ l) = 186 µM.

The concentration of GM130 in the cytoplasm was determined with the measured volume of the cytosol of HeLa cells of 0.94 x 10^-12^ l (Fujioka et al., 2006). The concentration of GM130 in the cytoplasm is c_GM130/cytoplasm_ = n_GM130/cell_ / V_cytoplasm_ = (1.43 x 10^-19^ mol) / (0.94 x 10^-12^ l) = 146 nM. The restricted localization of GM130 on the Golgi therefore increases the local concentration compared to the cytoplasm by 186 µM / 146 nM = 1.3 x 10^3^ fold.

## Key Reagents and Recourses

| REAGENT or RESOURCE | SOURCE | IDENTIFIER |
| --- | --- | --- |
| **Antibodies** |  |  |
| Mouse monoclonal anti-DDDDK (AP1501) | Bio X Cell | Cat# RT0269, RRID: AB_2687792 |
| Rabbit polyclonal anti-FLAG | Sigma | Cat# F7425, RRID: AB_439687 |
| Rabbit polyclonal anti-FXR1 | Sigma | Cat# HPA018246, RRID: AB_1849204 |
| Mouse monoclonal anti-FXR1 (6BG10) | Millipore | Cat# 05-1529, RRID: AB_1977197 |
| Rabbit polyclonal anti-G3BP1 | Proteintech | Cat# 13057-2-AP, RRID: AB_2232034 |
| Mouse monoclonal anti-G3BP1 | Proteintech | Cat# 66486-1-Ig, RRID: AB_2819031 |
| Mouse monoclonal anti-GAPDH (GA1R) | Invitrogen | Cat# MA5-15738, RRID: AB_10977387 |
| Rabbit polyclonal anti-GM130 (MLO7) | (Diao et al., 2003) | N/A |
| Rabbit polyclonal anti-GM130 (morsel), affinity purified | (Wei and Seemann, 2009b) |  |
| Rabbit polyclonal anti-GM130 (morsel) serum | (Wei and Seemann, 2009b) |  |
| Rabbit polyclonal anti-GM130 | Proteintech | Cat# 11308-1-AP, RRID: AB_2115327 |
| Mouse monoclonal anti-GM130 | BD Biosciences | Cat# 610822, RRID: AB_398141 |
| Rabbit polyclonal anti-Golgin-84 | (Beard et al., 2005) | N/A |
| Mouse monoclonal anti-Golgin-97 (CDF4) | Invitrogen | Cat# A-21270, RRID: AB_221447 |
| Rabbit polyclonal anti-GRASP55 | Proteintech | Cat# 10598-1-AP, RRID: AB_2113473 |
| Mouse monoclonal anti-GRASP55 | BD Biosciences | Cat# G34320-050 |
| Rabbit polyclonal anti-GRASP65 (UT465) | (Wei et al., 2015) | N/A |
| Mouse monoclonal anti-GRASP65 (D-12) | Santa Cruz | Cat# sc-374423, RRID: AB_10991322 |
| Rabbit polyclonal anti-p115 | Proteintech | Cat# 13509-1-AP, RRID: AB_2257094 |
| Mouse monoclonal anti-PABPC1 | Proteintech | Cat# 66809-1-Ig, RRID: AB_2882152 |
| Mouse monoclonal anti-TGN38 | BD Biosciences | Cat# T69020, RRID: AB_2315453 |
| Mouse monoclonal anti-alpha Tubulin (DM1A), eBioscience | Invitrogen | Cat# 14-4502-82, RRID: AB_1210456 |
| Alexa Fluor 488-conjugated highly cross-absorbed goat anti-mouse IgG (H+L) | Invitrogen | Cat# A-11029, RRID: AB_2534088 |
| Alexa Fluor 594-conjugated highly cross-absorbed goat anti-mouse IgG (H+L) | Invitrogen | Cat# A-11032, RRID: AB_2534091 |
| Alexa Fluor 647-conjugated highly cross-absorbed goat anti-mouse IgG (H+L) | Invitrogen | Cat# A-21235, RRID: AB_2535804 |
| Alexa Fluor 488-conjugated highly cross-absorbed goat anti-rabbit IgG (H+L) | Invitrogen | Cat# A-11034, RRID: AB_2576217 |
| Alexa Fluor 555-conjugated highly cross-absorbed goat anti-rabbit IgG (H+L) | Invitrogen | Cat# A-21428, RRID: AB_2535849 |
| Alexa Fluor 594-conjugated highly cross-absorbed goat anti-rabbit IgG (H+L) | Invitrogen | Cat# A-11037, RRID: AB_2534095 |
| HRP-conjugated goat anti-rabbit IgG | Jackson ImmunoResearch | Cat# 111-035-144, RRID: AB_2307391 |
| HRP-conjugated goat anti-mouse IgG | Invitrogen | Cat# 31430, RRID: AB_228307 |
| Purified rabbit IgG | ImmunoReagents | Cat# Rb-003-V |
| **Bacterial and Virus Strains** |  |  |
| BL21-Gold(DE3)pLysS E. coli | Aglient | Cat# 230134 |
| Rosetta 2(DE3) Singles Competent Cells | MilliporeSigma | Cat# 71-400-3 |
| **Chemicals, Peptides, and Recombinant Proteins** |  |  |
| Anti-FLAG M2 affinity gel | Sigma | Cat# A2220 |
| cOmplete protease inhibitor cocktail | Roche | Cat# 11873580001 |
| DiD perchlorate | AAT Bioquest | Cat# 22034 |
| DSS (disuccinimidyl suberate) | Thermo Scientific | Cat# A39267 |
| Dynabeads Oligo (dT)<sub>25</sub> | Invitrogen | Cat# 61002 |
| Dynabeads Protein A | Invitrogen | Cat# 10001D |
| 3x FLAG peptide | Apexbio | Cat# A60014 |
| Fluorescein Dextran, 70 kDa, Lysin fixable | Invitrogen | Cat# D1822 |
| 1,6-Hexanediol | Sigma-Aldrich | Cat# 240117 |
| HIS Lite OG488-Tris NTA-Ni complex | AAT Bioquest | Cat# 12615 |
| Hoechst 33342 | Invitrogen | Cat# H21492 |
| IPTG | Fisher | Cat# BP1620-1 |
| 3-Indole-acetic acid sodium salt (IAA) | Abcam | Cat# ab146403 |
| Lipofectamine RNAiMAX | Invitrogen | Cat# 13778075 |
| Ni-NTA Agarose | Qiagen | Cat# 30210 |
| PEI Max | Polysciences | Cat# 24765 |
| Protein A/G PLUS agarose | Santa Cruz | Cat# sc-2003 |
| RNasin Plus Ribonuclease Inhibitor | Promega | Cat# N2615 |
| RNase A | Sigma-Aldrich | Cat# R5125 |
| Sodium arsenite | Sigma-Aldrich | Cat# S7400 |
| Strep-Tactin XT 4Flow resin | IBA Lifesciences | Cat# 2-5010-010 |
| SDS | Fisher | Cat# BP166-500 |
| Texas Red Dextran, 70 kDa, Lysin fixable | Invitrogen | Cat# D1864 |
| **Critical Commercial Assays** |  |  |
| Direct-zol RNA miniprep kit | Zymo Research | Cat# R2050 |
| Duolink In Situ Red starter kit<br>Mouse/Rabbit | Sigma | Cat# DUO92101 |
| **Experimental Models: Cell Lines** |  |  |
| HEK 293T | ATCC |  |
| Normal rat kidney (NRK) | ATCC |  |
| RC55 | (Zhang and Seemann, 2021) |  |
| RC65 | (Zhang and Seemann, 2021) |  |
| SV589 (immortalized human fibroblasts) | (Yamamoto et al., 1984) |  |
| **Oligonucleotides** |  |  |
| For oligoes, see Table S |  |  |
| **Recombinant DNA** |  |  |
| pLVX-NAGTI-GFP-Puro | (Zhang and Seemann, 2021) | N/A |
| pcDNA3.1(-) GM130 | (Wei et al., 2015) | N/A |
| pcDNA3.1(-) GM130 FL (long version) | This paper | N/A |
| pEGFP-N1 GM130 FL (long version) | This paper | N/A |
| pET23a(+) GM130 N74-GST | (Wei et al., 2015) | N/A |
| pET23a(+) GM130 N74-his | (Wei et al., 2015) | N/A |
| pET23a(+) GM130 N74-Strep | This paper | N/A |
| pET23a(+) GM130 N122-his | This paper | N/A |
| pET23a(+) GM130 N122-oDi-his | This paper | N/A |
| pET23a(+) GM130 N122-oTet-his | This paper | N/A |
| pEGFP-N1 vector | Clontech | Cat# 6085-1 |
| pSMPP-mCherry-hTRIM21 | (Clift et al., 2017) | Addgene #104972 |
| psPAX |  | Addgene |
| pVSVG |  | Addgene |
| SpyCatcher002-oDi | (Khairil Anuar et al., 2019) | Addgene #124661 |
| SpyCatcher002-oTet | (Khairil Anuar et al., 2019) | Addgene #124663 |
| **Software and Algorithms** |  |  |
| Fiji (ImageJ2) | Fiji | https://fiji.sc/ |
| Prism 9 software | GraphPad | https://www.graphpad.com |

## Oligonucleotides

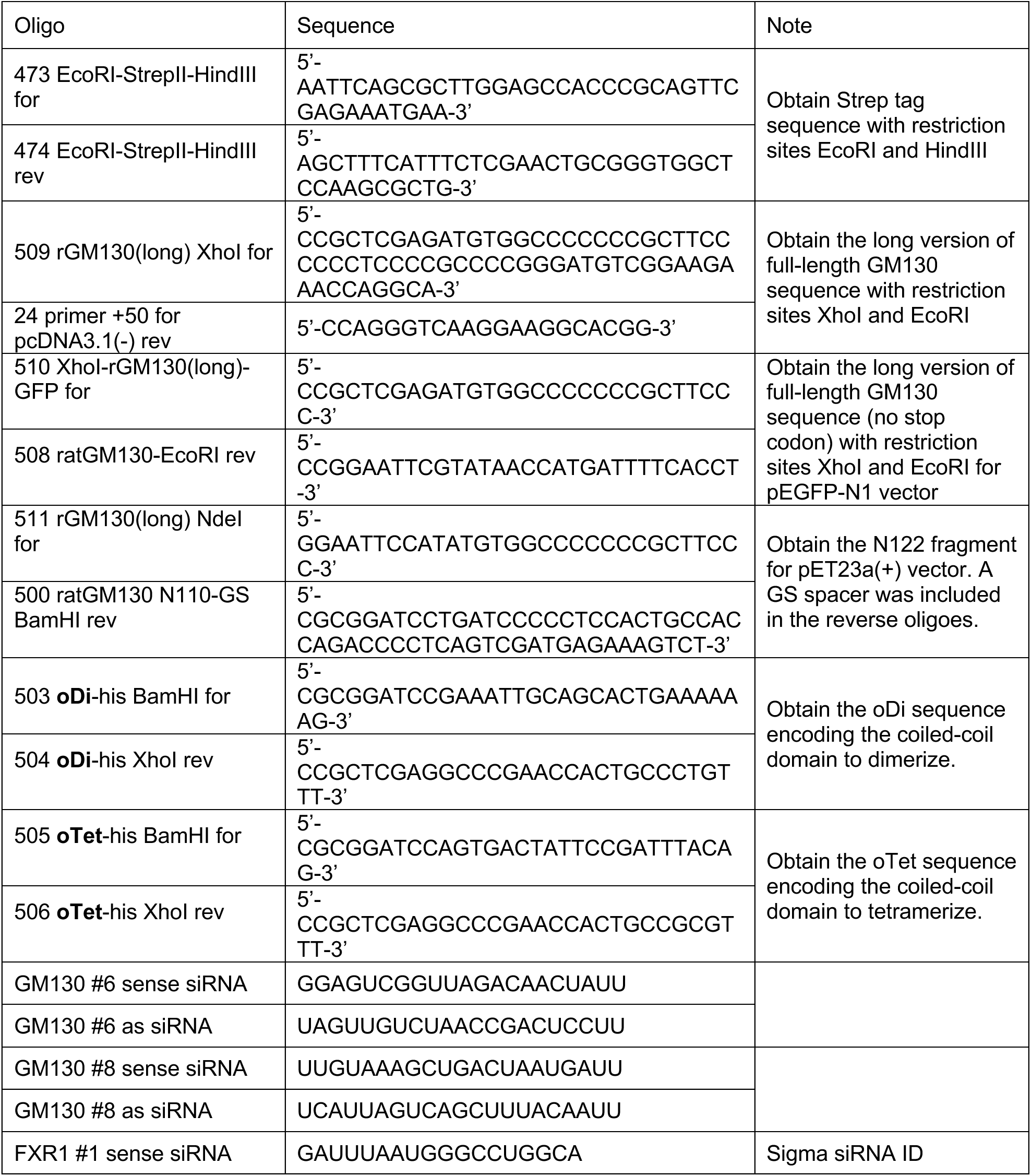
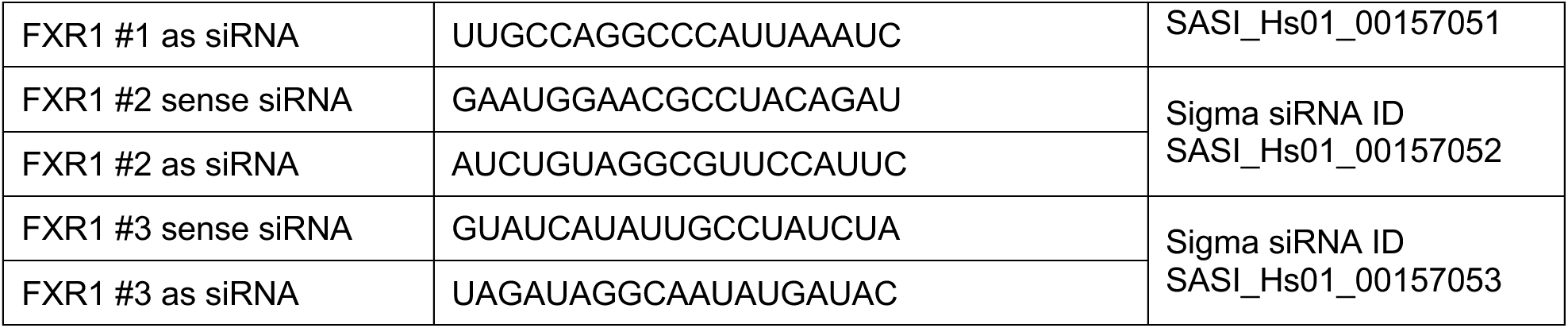

## Acknowledgments

We thank Jen-Hsuan Wei and Haijing Guo for suggestions and insightful discussions, the Proteomics Core Facility for mass spectrometry analysis, and the Live Cell Imaging facility at the University of Texas Southwestern Medical Center for imaging support. This work was supported by grants from the NIH (GM096070) and the Welch Foundation (I-1910).

## Author contributions

Y.Z. and J.S. designed the project, performed the experiments, analyzed the data, prepared the figures and wrote the manuscript.

---

**Fig.S1.**
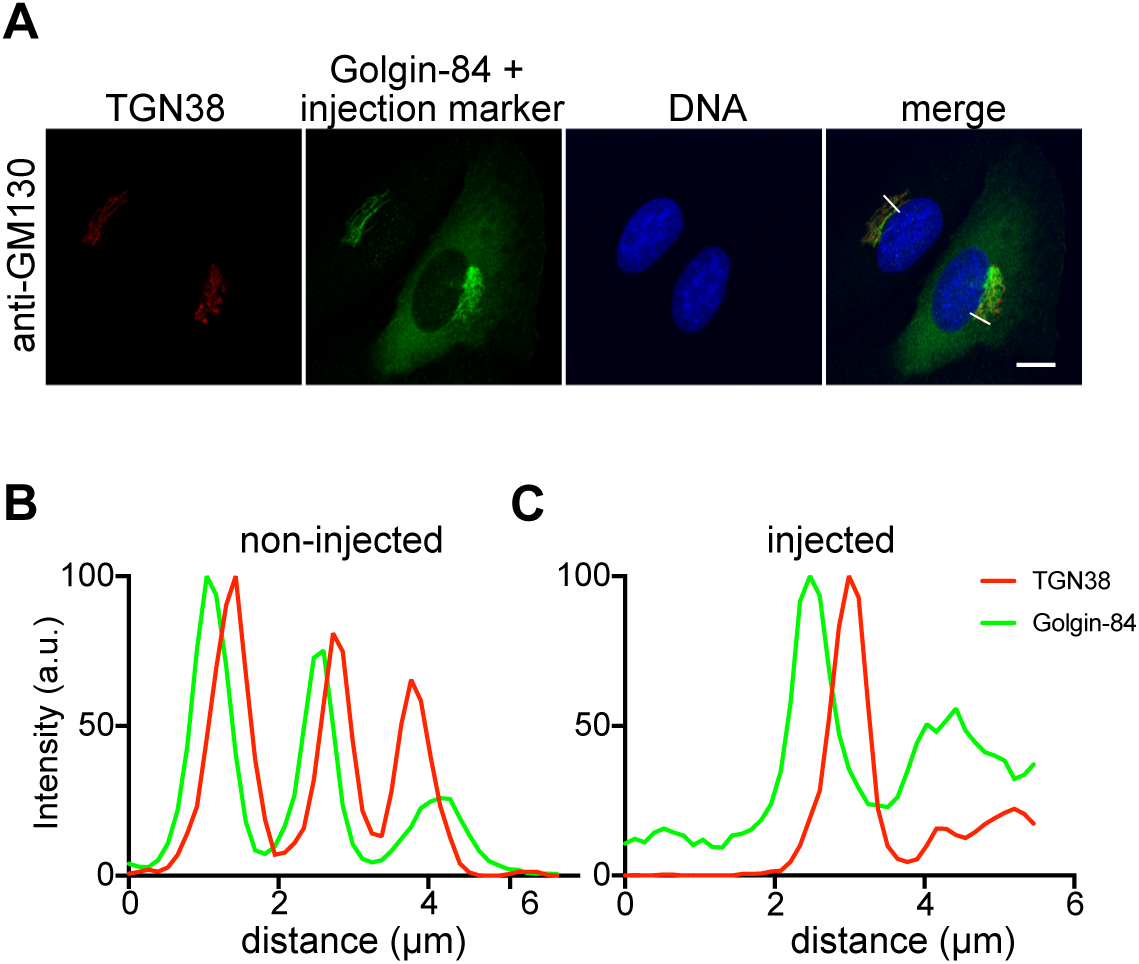
Acute degradation of GM130 does not affect Golgi stacking. (A-C). NRK cells stably expressing mCherry-TRIM21 were injected with anti-GM130 together with fluorescent dextran as an injection marker. Cells were fixed 2 hours later and immunostained for TGN38 (red) to mark the TGN, Golgin-84 resident to the cis-Golgi (green) and labeled for DNA (blue). White lines on top left non-injected cell and injected cells (bottom) indicate the line scan of the fluorescence intensities shown in (B) and (C), respectively. Scale bar, 10 µm.

**Fig.S2.**
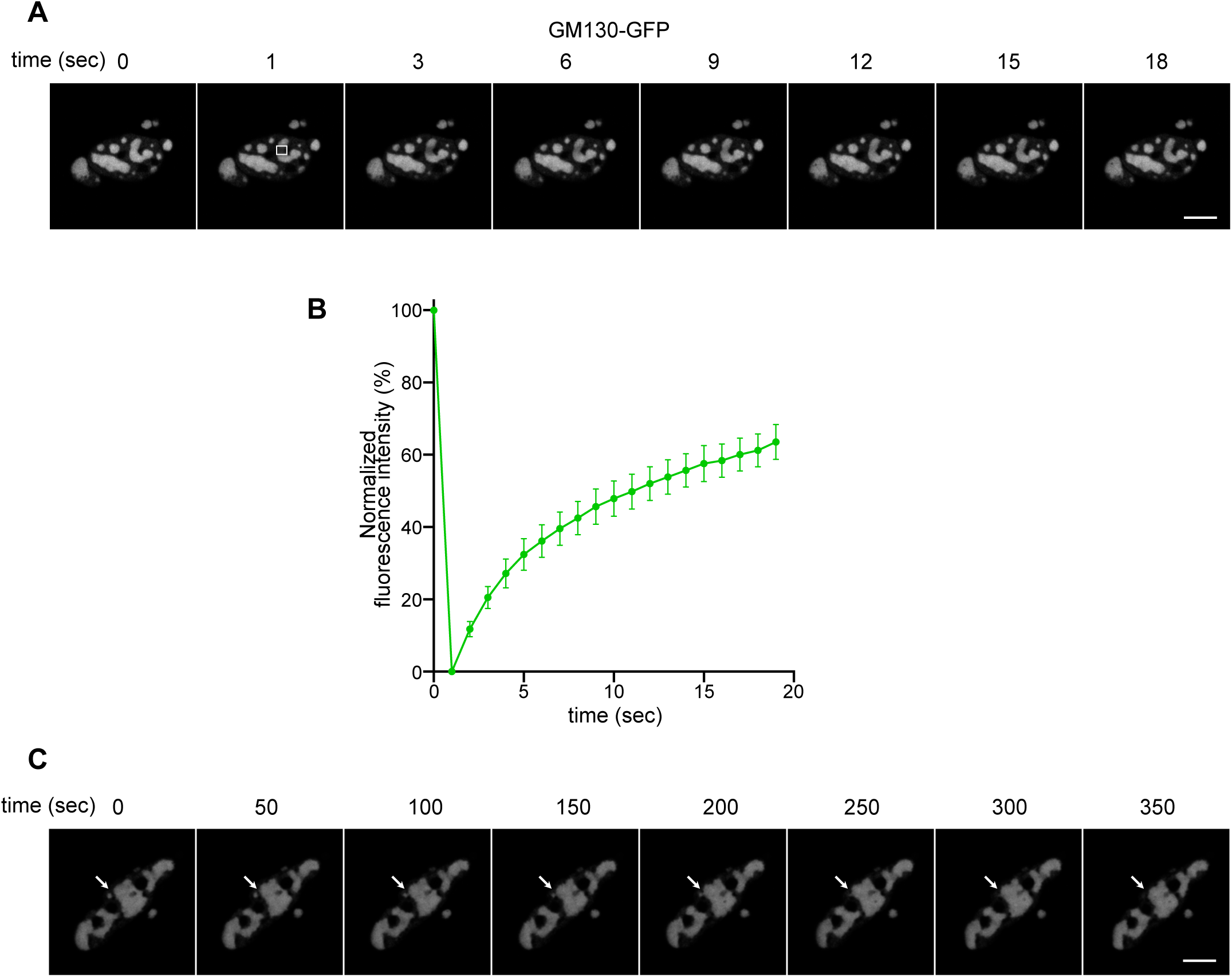
GM130 condensates in cells are highly dynamic. (A). GM130-GFP was transiently overexpressed and condensates were subjected to FRAP analysis. Representative images are shown at the indicated time points, with white box marking the photobleached area of the Golgi. Scale bar, 10 µm. (B). Quantitation of the FRAP results. The recovery rate at each time point was calculated as the ratio of the mean intensity of the photobleached area to that of the adjacent area and then normalized to the closest time point before bleaching. n = 18. Error bars represent mean ± SEM. (C). Representative time lapse images of GM130-GFP at indicated time points show the fusion of GM130 condensates (arrow heads). Scale bar, 10 µm.

**Fig. S3.**
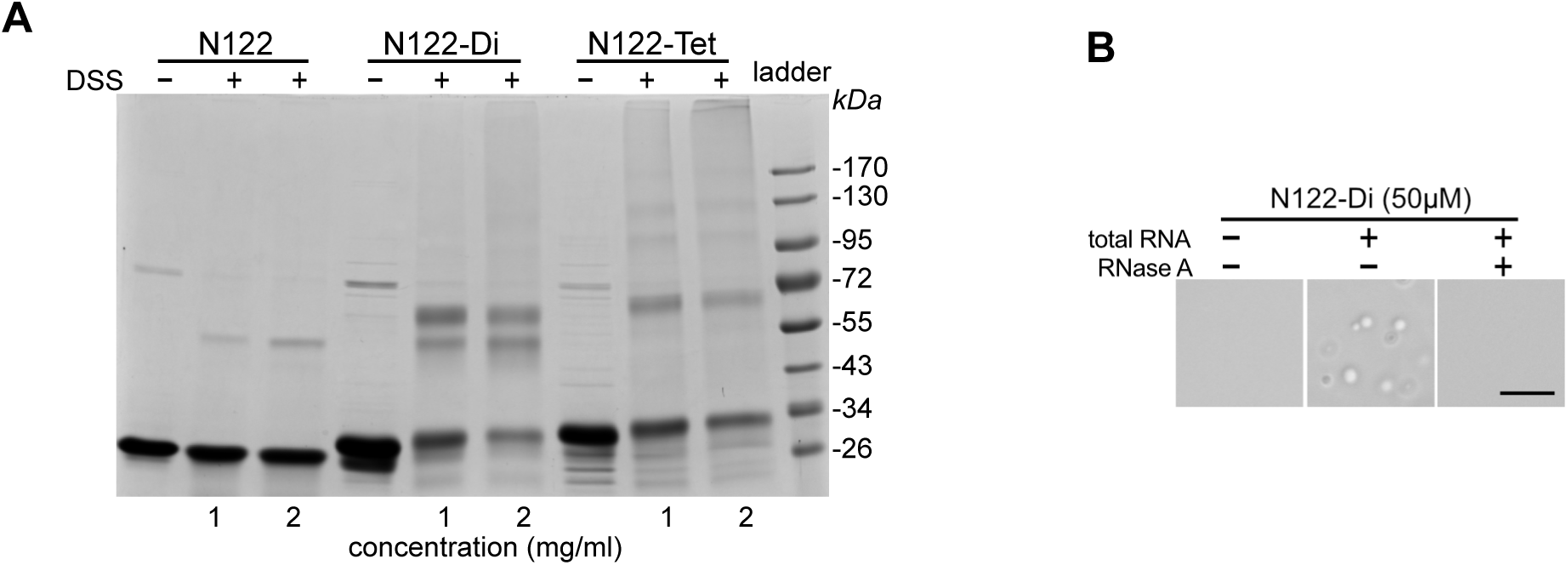
(A). GM130 N122, N122-Di and N122-Tet at indicated concentrations were incubated with the cross- linker disuccinimidyl suberate (DSS) and then analyzed by SDS-PAGE and Coomassie blue staining. (C). RNase A prevents LLPS of N122-Di and total RNA. LLPS was performed with N122-Di and 100 ng/µl total RNA, with or without the addition of 1mg/ml RNase A. DIC images of droplets were captured within 5 min after LLPS induction. Scale bar, 10 µm.

**Fig. S4.**
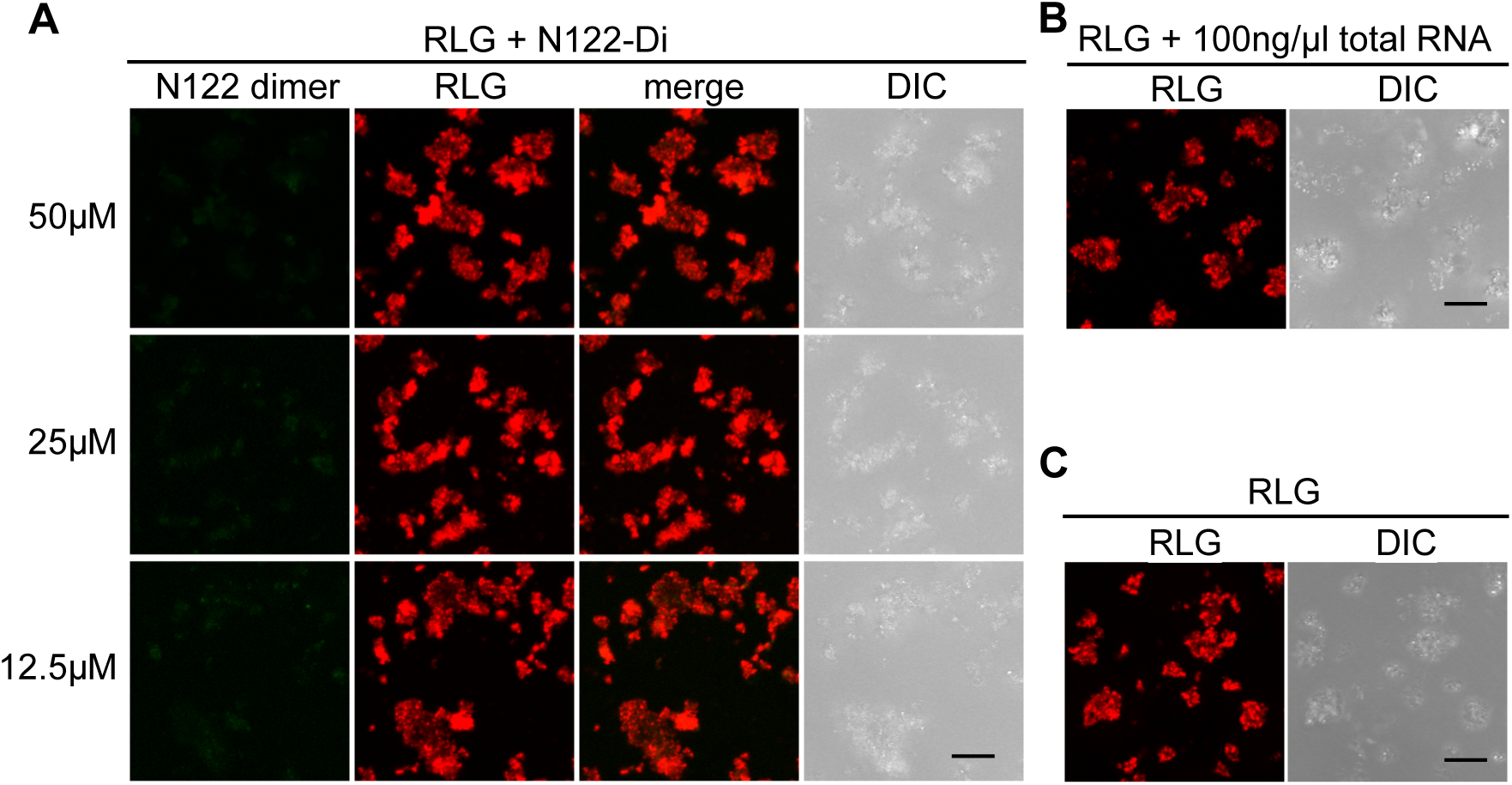
Condensates formed by GM130 N122-Di and RNA associate with Golgi membranes in vitro. (A). LLPS was assessed by mixing fluorescently labelled rat liver Golgi membranes (RLG) with N122-Di (containing 5% fluor 488-labeled N122-Di) at different concentrations without addition of RNA. Images were taken 30min later to allow the sedimentation of RLG. Scale bar, 10 µm. (B-C). Controls for (A) were set by mixing rat liver Golgi and 100 ng/µl total RNA from SV589 cells (B) or by mixing rat liver Golgi with regular salt buffer only. Images were taken 30min later to allow the sedimentation. Scale bar, 10 µm.

